# Language model generates *cis-*regulatory elements across prokaryotes

**DOI:** 10.1101/2024.11.07.622410

**Authors:** Yan Xia, Jinyuan Sun, Xiaowen Du, Zeyu Liang, Wenyu Shi, Shuyuan Guo, Yi-Xin Huo

## Abstract

Deep learning had succeeded in designing *Cis*-regulatory elements (CREs) for certain species, but necessitated training data derived from experiments. Here, we present Promoter-Factory, a protocol that leverages language models (LM) to design CREs for prokaryotes without experimental prior. Millions of sequences were drawn from thousands of prokaryotic genomes to train a suite of language models, named PromoGen2, and achieved the highest zero-shot promoter strength prediction accuracy among tested LMs. Artificial CREs designed with Promoter-Factory achieved a 100% success rate to express gene in *Escherichia coli*, *Bacillus subtilis*, and *Bacillus licheniformis*. Furthermore, most of the promoters designed targeting *Jejubacter* sp. L23, a halophilic bacterium without available CREs, were active and successfully drove lycopene overproduction. The generation of 2 million putative promoters across 1,757 prokaryotic genera, along with the Promoter-Factory protocol, will significantly expand the sequence space and facilitate the development of an extensive repertoire of prokaryotic CREs.

## Introduction

Transcription, a pivotal process in the central dogma, is primarily regulated by CREs, especially promoters^1,2,3^. In biological research, promoters serve as indispensable tools for studying gene expression and regulation^4,5^. Customized promoters are also widely used in biotechnology applications to enable controlled gene expression, developing biosensors for molecules of interests, and fine-tuning metabolic flux in pathways^6,7,8,9,10^. However, only a limited number of promoters, primarily from a few model organisms, have been well-characterized and engineered as bioparts for research and applications^11^. Thus, the range of achievable gene expression levels are restricted by the narrow repertoire^12^. Also, repetitively introduction of these endogenous elements may lead to genetic instability and disruption of native cellular processes in host organisms^13^. Beyond those limitations, a greater challenge is the lack of identified promoters for non-model organisms^14^. The limited availability of promoters for non-model organisms has hindered the development of new chassis that offer various advantages over traditional model organisms.

To expand the spectrum of promoters for desired chassis, mining exogenous promoters and engineering known promoters were emerged as two primary approaches^12,15,16^. Due to the relation between DNA sequences and potential regulatory functions remained elusive, both the mining and engineering relied on experimental characterizations of sequences derived from synthesized oligos or random mutagenesis^17,18^. Such processes presented challenges in terms of time and resource requirements therefore cannot be applied across the broader prokaryotic kingdom^19^. Consequently, there is an urgent need for more efficient and broadly applicable methods to enhance the availability promoters to enable controlled gene transcription in desired organisms.

Recently, machine learning based methods have shown promise in designing promoters by leveraging experimentally derived datasets^20,21,22,23^. These approaches offered a solution for expanding promoter repositories in well-studied model organisms and optimizing the expression level on demands^24^. However, the requirement of labeled datasets presented great challenge to develop a universal method for regulator design with unlimited species scope^24,25^. From a machine learning perspective, design promoters for various species are different tasks, and for the most cases the task-specific training data are limited. As the Generative Pre-trained Transformers (GPTs) was proved as few shot learners^26^, training a foundation model on a large number of CREs can be a possible solution to enable the model to generalize to underrepresented or even previous unseen species. We previously trained the PromoGen model on experimentally derived prokaryotic promoter sequences and successfully generated artificial *Bacillus subtills* promoters upon fine-tuning^27^. Building upon these insights, we sought to expand the scope and capabilities of this pre-training and fine-tuning paradigm to develop a universal solution for designing promoters across a wide range of prokaryotes.

In this study, we present the PromoGen2 language model series, trained on a newly curated dataset of 17 thousand prokaryotic genomes. PromoGen2 showed improved zero-shot promoter activity prediction over other nucleotide language models and significantly better generation quality than PromoGen when fine-tuned on the same data. Experimental validation in *B. subtilis*, a species with stringent promoter requirements, revealed that all generated promoters were active, with 62% surpassing the native strong promoter *P_43_*. Additionally, we designed promoters for *Jejubacter sp.* L23, achieving 95% active ratio after fine-tuning. Using lycopene production as an example, we demonstrated that generated promoters could regulate yield over a 3.68-fold range. Furthermore, we introduced a conditional generative model named PromoGen2-proka to produce 2 million genus-level promoter sequences covering 1,757 genera. PromoGen2 thus offers a universal tool for *de novo* promoter generation in prokaryotes, advancing synthetic biology.

## Results

### Nucleotide language model trained on expanded dataset of CREs

In scenarios where domain-specific data is limited, specific tasks were performed by fine-tuning large language models (LLMs) trained on vast corpora as evidenced in the BERT^28^ and GPT^26^. To maximize the knowledge of LLMs about CREs, we initially collected 59.9 million upstream untranslated region (UTR) sequences containing the 160 nucleotides upstream of each coding sequence (CDS) from 17,806 NCBI prokaryotic genomes (**Fig. 1a**), to construct the most comprehensive dataset of prokaryotic upstream regulatory sequences to date, named prokaryotic untranslated region dataset (PURD). The data in PURD were clustered and scored by convolutional neural network (CNN) model, and the 1.4 million sequences with highest scores were collected and named PURD-core, which is ten times larger than the existing Prokaryotic Promoter Database (PPD) **(Supplementary Fig. 1)**. PURD-core was served as the pre-training dataset for PromoGen2, a decoder-only Transformer (**Fig. 1b**) with the next-token prediction objective (**Fig. 1c**). The nucleotide sequences were tokenized (split into a word) on single nucleotide resolution **(Supplementary Fig. 2)**. We processed all training data in both 5’-to-3’ and 3’-to-5’ orientations to mitigate potential information loss given that GPT model is an autoregressive model inherently capturing only the dependency of downstream nucleic acid sequences on upstream sequences (**Fig. 1d**). We trained three PromoGen2 models including 8M, 33M and 149M parameters, respectively, and evaluated the ability of these models to capture the distribution of native promoter sequences by assessing their perplexity on unseen sequences (**Fig. 1e**). The perplexity scores obtained for PromoGen2-xsmall, -small, and -base models on our test dataset were 1.195, 1.191, and 1.186, respectively, demonstrating that larger PromoGen2 models obtained reduced perplexity, enhancing feature acquisition without succumbing to overfitting tendencies.

**Fig. 1.**
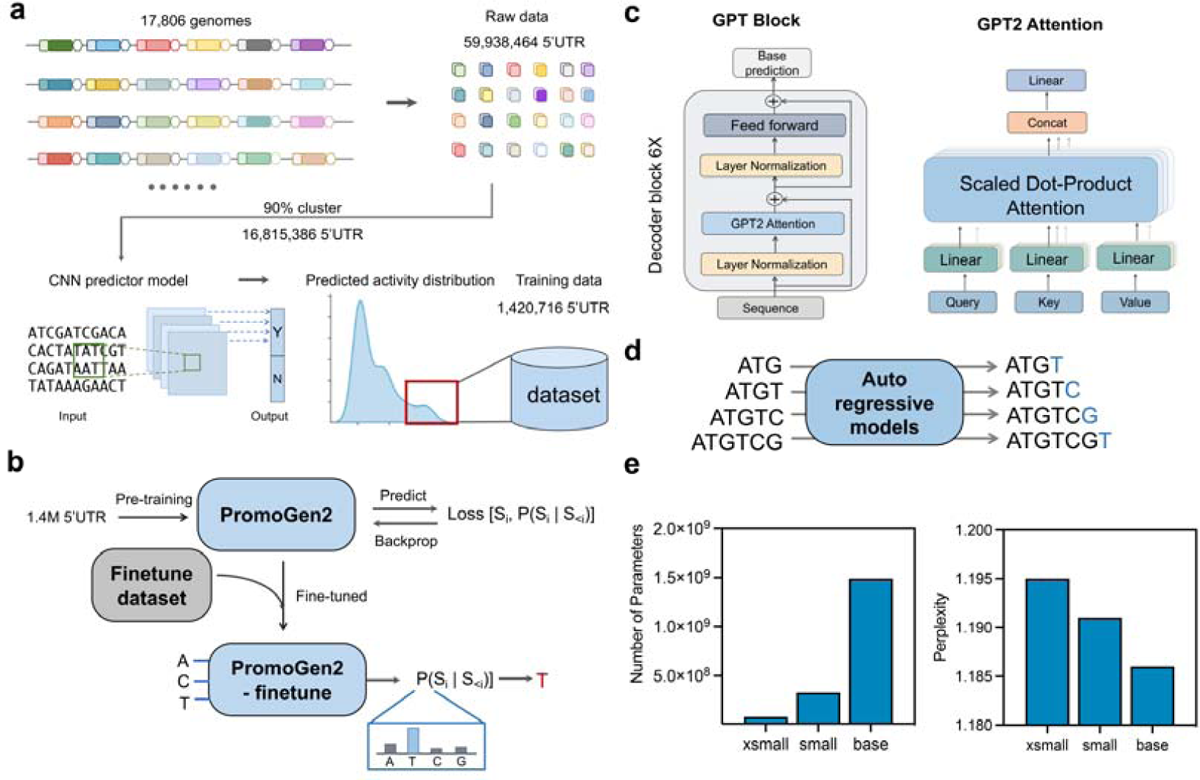
Overview of PromoGen2 models for prokaryotes promoter sequence generation. **a.** Training Dataset Process in PromoGen2 Models. The PromoGen2 models utilize a comprehensive training dataset, which consists of diverse 5’ upstream untranslated regions (UTRs) from various prokaryotic species. **b.** Diagram of the model training. The training process of PromoGen2 involves two key stages: an initial pre-training phase and a subsequent fine-tuning phase. During pre-training, the model is exposed to a broad spectrum of 5’ UTR sequences from multiple prokaryotic species. In the fine-tuning phase, the model is specifically adapted to generate promoter sequences for particular species by focusing on their unique 5’ UTRs or promoters. **c.** The Transformer architecture of PromoGen2. The self-attention mechanism is crucial for capturing and understanding the distribution patterns within the 5’ UTR sequences. **d.** Diagram of promoter generation in an autoregressive manner. This means that the model generates each nucleotide of the promoter sequence step-by-step, using the previously generated nucleotides as context. **e.** The parameters and perplexity of the PromoGen2 pretrain models.

### Generative modeling captured essential features of promoters

To evaluate the diversity of model-generated sequences, we constructed a phylogenetic tree using 10,000 sequences uniformly sampled from PURD-core and 10,000 generated sequences by each PromoGen2 model. **Fig. 2a and Supplementary Fig. 3** demonstrated that all three PromoGen2 models could generate diverse sequences that effectively cover the space of native promoters. To evaluate the effects of sampling strategy, *i.e.* the hyperparameter settings of temperature (T) and nucleus sampling (P) values, on the characteristics of sequences generated by PromoGen2-xsmall, -small, and -base, 10,000 sequences were generated individually under different settings, *i.e.* 1 or 0.9. Results demonstrated that lower T and P values increased typical motifs and predicted activity (**Fig. 2b**) and decreased the GC content (**Fig. 2c**) in the expense of introducing longer homopolymers (**Fig. 2d**), in agreement with the well-known effects of the sampling strategy and general-believed trade-off between diversity and quality of the generated sequences^29,30^. Taken together, sampling strategy plays a bigger role than model size on generating sequencing diversity, which could reduce the occurrence and average length of DNA repeats, the main cause of the synthesis difficulty and the instability of artificially synthesized DNA. Totally 60 thousands of sequences were generated using two sampling strategies by the three models and all of them were predicted to be active by our CNN scoring model^27^(**Fig. 2e**), while the distribution of predicted scores of the generated promoters were shifted toward higher score when T and P were lowered to reduce the sequence randomness or diversity.

**Fig. 2.**
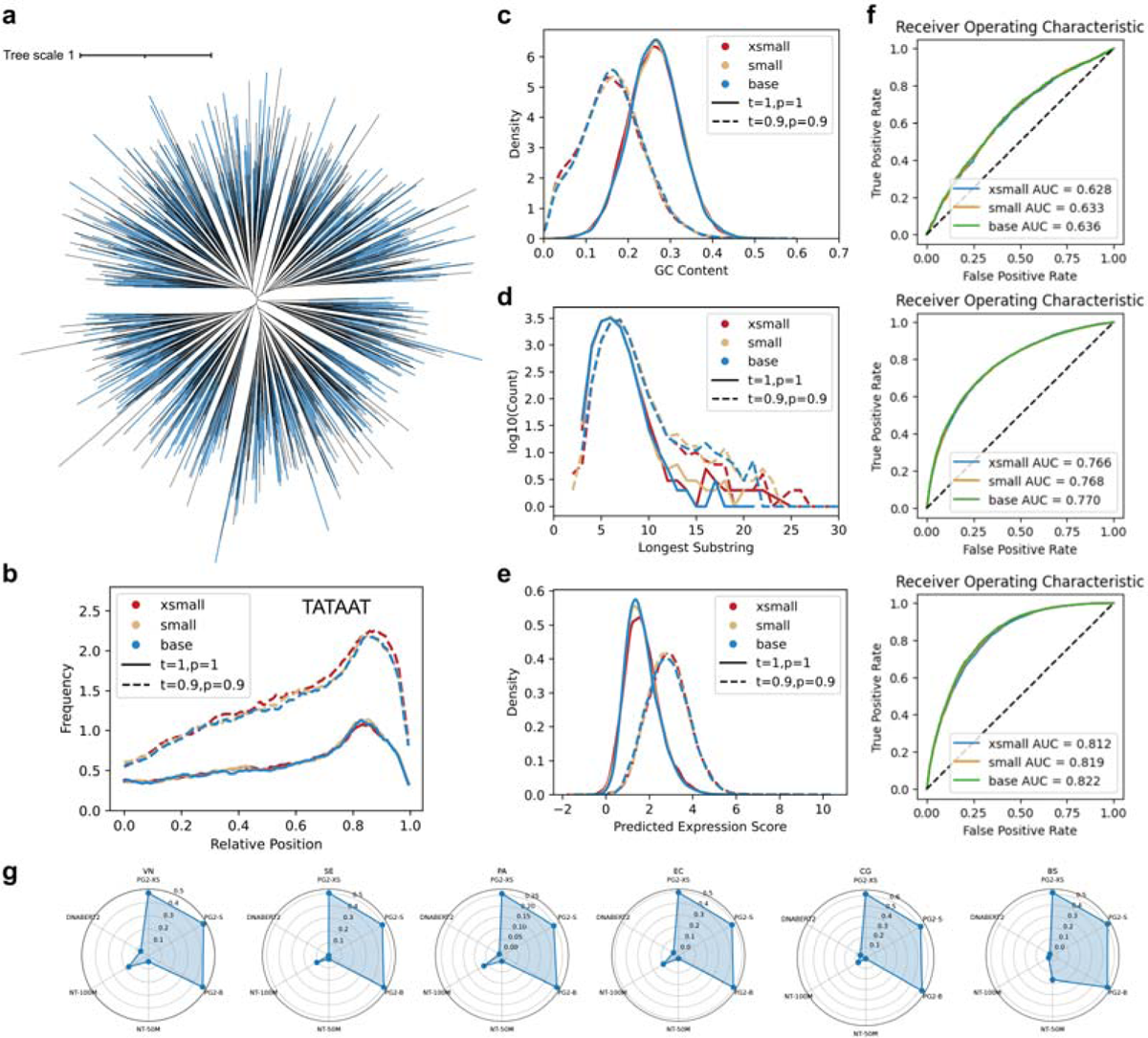
The evaluation of pre-training PromoGen2 in various species. **a.** The Phylogenetic tree of PromoGen2-base generated sequence (blue) and native promoters (black)**. b.** Analysis of DNA shape for generated sequences and random sample sequences at the TATAAT motif using PromoGen2-xsmall, small, and base. **c.** The GC content of generated sequence by PromoGen2-xsmall, small, and base. **d.** Consecutive identical nucleotides in the generated sequences. **e.** The prediction of generated sequence activities by CNN. **f.** Evaluation of the receiver operating characteristic (ROC) curves for the three pre-trained PromoGen2 models on *B. subtilis*, *E. coli,* and *P. aeruginosa*. **g.** Scoring the predictive capabilities of PromoGen2-xsmall, PromoGen2-small, PromoGen2-base, Nucleotide Transformer (N.T) −50, N.T-100, and DNABERT2 on promoters from *E. coli*, *B. subtilis*, *S. enterica*, *V. natriegens*, *C. glutamicum,* and *P. aeruginosa* in translational levels.

Predicted promoters were classified by PromoGen2 from a weak-selection-pressure dataset for three individual species, and the Area Under the Receiver Operating Characteristic curve (AUROC) were used to compute the classification accuracy, which was generally increased along with our model size (**Fig. 2f**). For example, the AUROC for *P. aeruginosa* dataset achieved 0.82 by PromoGen2-base, indicating good discriminative ability. However, the middle-sized model (33M parameters) occasionally exhibited inferior performance compared to the smallest models in the ranking task **(Supplementary Fig. 4)**.

In some datasets, the promoter activities were tested and characterized by the expression level of the GFP, assuming that the regulation does not occur in the translational level. The Spearman correlation coefficients between tested and zero-shot predicted promoter activities were calculated for six selected species from a GFP-based dataset and the average numbers were 0.475, 0.472, and 0.481 for PromoGen2-xsmall, -small, and -base, respectively (**Fig. 2g**), while other nucleotide language models with similar parameter size, *i.e.* Nucleotide Transformer (50M and 100M versions) and DNABERT2 (117M), failed **(Supplementary Table 1)**, demonstrating that preset data filtration was needed for zero-shot promoter strength predictions by PromoGen2.

### Language model enabled generally applicable promoter design for prokaryotes

The PromoGen2-xsmall is of identical architecture and parameter size as the PromoGen, and both were fine-tuned using PPD data of 28 main species. This provided a fair comparison of the effect of the pre-training data, which is the only difference between PromoGen2-xsmall and PromoGen. Also, we tested the conditional generation schema using species-level control tag to prompt a fine-tuned species-specific generation model **(Supplementary Fig. 5)**. We employed three distinct fine-tuning methods for multi-species generation including: 1) we fine-tuned a model for each species separately without species labels to produce “PromoGen2-xsmall single-no tag” as in our previous study^32^, 2) we explored single-species level conditional generation by fine-tuning with the PPD dataset tagged with species labels to create “PromoGen2-xsmall single-tagged”, 3) we fine-tuned PromoGen2-xsmall with all species within the PPD dataset to obtain “PromoGen2-xsmall PPD” (**Fig. 3a**).

**Fig. 3.**
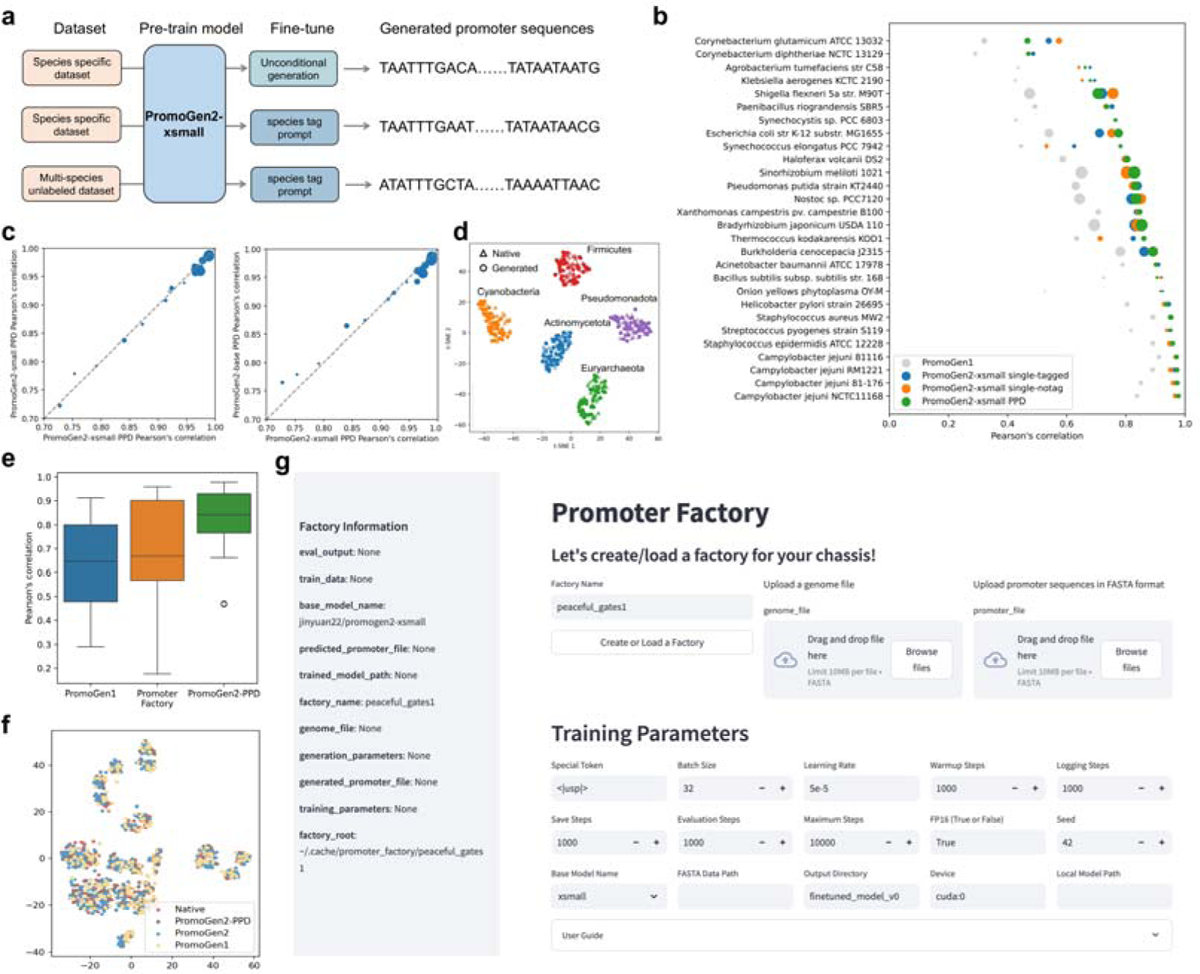
The PromoGen2 captures key characteristics across multiple species. **a.** Diagram illustrating the fine-tuning of the PromoGen2-xsmall model using three different methods. **b.** Pearson’s r for the *de novo* generated promoters of 27 major species in the PPD. Different circle sizes represent varying data volumes. **c.** A 2D projection plot depicting the distribution of native promoters and sequences generated by PromoGen2. The 28 species belong to five classes: Firmicutes (red), Cyanobacteria (yellow), Pseudomonadota (purple), Actinomycetota (blue), and Eurarchaetoa (green). **d.** Comparison of the Pearson’s r for promoters generated by PromoGen2-xsmall, PromoGen2-small, and PromoGen2-base across 27 species. **e.** The evaluation of generated sequence quality of PromoGen1, Promoter-Factory and PromoGen2-PPD. **f.** A 2D projection plot illustrating the distribution of native promoters alongside sequences generated by PromoGen1, Promoter-Factory, and PromoGen2-PPD. **g.** GUI of the Promoter-Factory. This involves using Prodigal to predict CDSs, extracting upstream regions, applying the Promoter Calculator for 60 bp core promoters, and fine-tuning the pre-trained PromoGen2 model. The fine-tuned model generates 60 nt promoters for each species, ranking them by score, with lower scores indicating higher promoter likelihood. Users can upload a genome, generate species-specific promoters, and adjust hyperparameters including batch size, learning rate, warmup steps, logging steps and so on. PromoGen2 also supports direct promoter sequence uploads without dataset processing.

As shown in **Fig. 3b and Supplementary Table 2**, the quality of promoters generated by PromoGen2-xsmall was overwhelmingly better than the one generated by PromoGen. Specifically, the Pearson’s r exceeded 0.5 for 26 out of 28 species and surpassed 0.9 for species such as *Campylobacter jejuni*, *Staphylococcus aureus*, *Helicobacter pylori*, and *B. subtilis*, indicating that fine-tuning enhances the quality of generated promoters. The averaged Pearson’s r of single- and multi-species conditional generation schema was improved to 0.82 and 0.83 (**Supplementary Fig. 6**), respectively, possibly due to that the learned embedding vector of additional control token can store species-level information. During the inference stage, the control tags will be used in the prompt and provide species-level information explicitly. For multi-species fine-tuned model, the generation quality will benefit further from the multi-tasking during training stage. The 28-species sequence logo analysis indicated that the sequence distribution generated by PromoGen2-xsmall PPD was entirely consistent with that of native sequences (**Supplementary Fig. 7**). Besides fine-tuning strategy, we also explored the effects of pre-trained model size on the sequence quality generated after fine-tuning and concluded that model size has positive effect on the sequence quality, especially in species with smaller fine-tuning datasets (**Fig. 3c**). This suggests that larger models are likely able to capture more diverse and deeper promoter patterns, thereby exhibiting superior few-shot generalization capabilities. The 28 major species in the PPD database belong to five phyla including Firmicutes, Pseudomonadota, Cyanobacteria, Actinomycetota, and Euryarchaeota. We prompted PromoGen2-xsmall single-tagged generated phylum-specific promoters, which clustered closely within the same phylum and separated from other phyla on a two-dimensional plane (**Fig. 3d**).

Considering the limited availability of experimentally verified promoters, we developed a computation protocol, named Promoter-Factory, to annotate promoters from genomic sequences and fine-tune the PromoGen2 subsequently in order to design CREs for targeting species. We tested this protocol on 28 species in the PPD. The results indicated that although the annotated promoters differ significantly from true promoters, they still guide the model’s generation distribution toward the promoters of the corresponding species when used as fine-tuning data (**Fig. 3e**). Furthermore, the Pearson’s r of the 6-mer frequency between the generated promoter sequences and native sequences reached 0.69, showing a significant improvement compared to fine-tuning directly on PromoGen and approaching the performance of fine-tuning with PPD experimental data (**Fig. 3e**). We projected native promoters, PromoGen2-PPD, Promoter-Factory, and PromoGen-generated promoters onto a two-dimensional plane, showing species-specific clustering and even distribution for different species (**Fig. 3f**). Our method can generate high-quality promoters from noisy genomic data without experimental input, with potential for application to any prokaryotic organism. We also developed a user-friendly graphical user interface (GUI) to enable non-expert to use Promoter-Factory easily (**Fig. 3g**).

### Evaluation the CREs designed with Promoter-Factory *in vivo*

We first validate Promoter-Factory to design CREs for *B. subtilis*, a gram-positive bacterium commonly used for protein expression and chemical production but limited by scarce promoter data^31^. We fed the Promoter-Factory with experimentally characterized promoters of *B. subtilis* to fine-tune PromoGen2-small resulted in the PromoGen2-bsu model (**Fig. 4a**). Sampling 50 sequences from PromoGen2-bsu, we performed wet-lab validation with *P_43_*and a random sequence as positive and negative controls, respectively (**Supplementary Table 3**). The results showed that all generated sequences have promoter activity, with 62% exhibiting higher expression than *P_43_* and promoters A1, A16, A17, and A20 have activities 2.32, 2.42, 2.44, and 2.64 times that of *P_43_* (**Fig. 4b**). Compared to our previous species-specific promoter generation model PromoGen-bsu^27^, the overall strength of promoters generated by PromoGen2 increased by 5.06-fold (**Fig. 4c**), indicating that PromoGen2, pre-trained with extensive data, better learned the universal characteristics of promoters, including the features of the flanking region of −10 and −35 regions. We conducted PWM analysis on 500 sequences generated by PromoGen2-bsu, and the results indicated that their features were largely consistent with those of native promoters and sequences generated by PromoGen-bsu **(Supplementary Fig. 8)**. These results indicated that suggests that the flanking regions of the −10 and −35 regions, despite the lack of significant sequence preference, plays an important role in promoter activity.

**Fig. 4.**
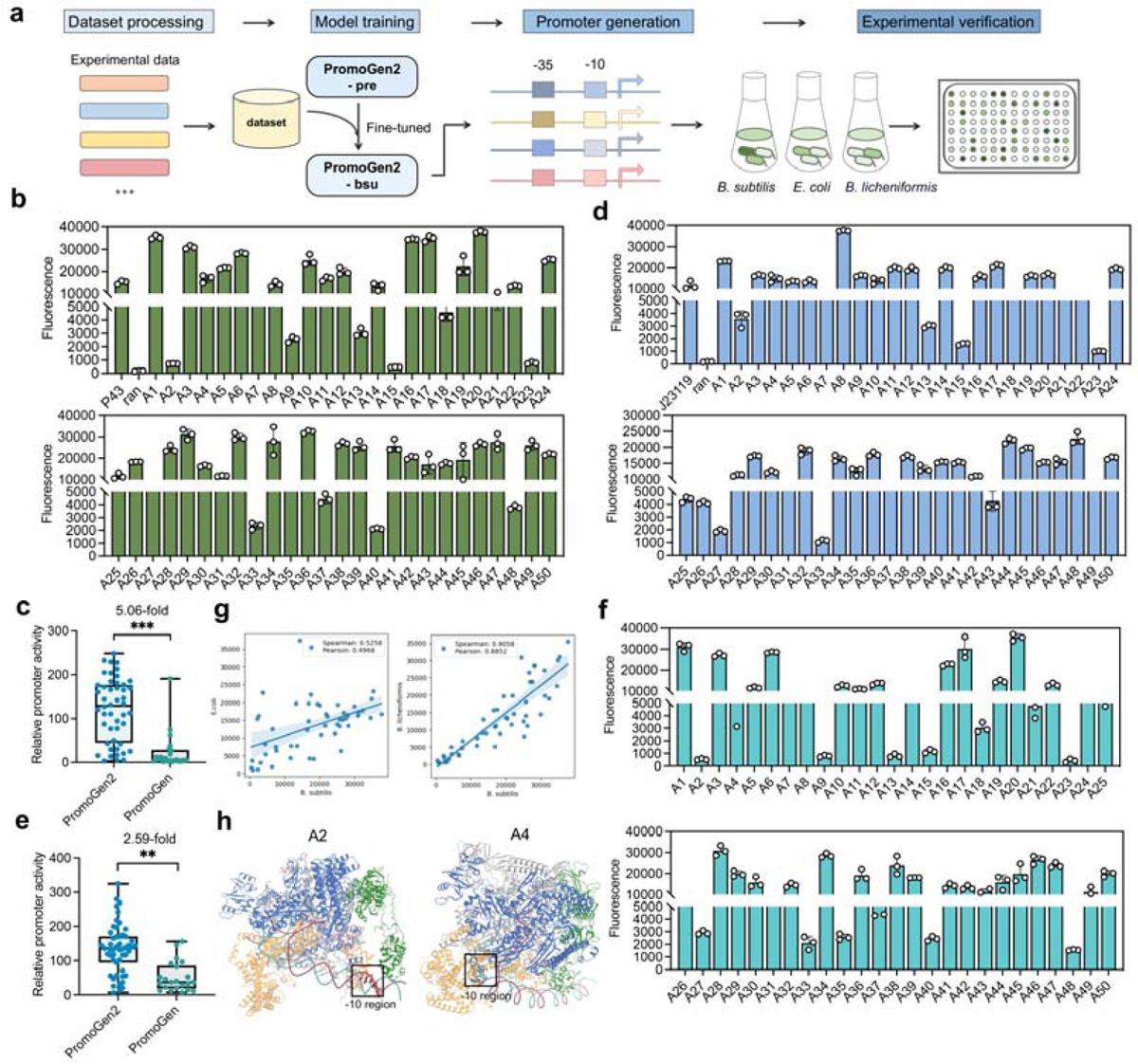
Validation of the generated promoter activities by PromoGen2-bsu in various species. **a.** Detailed description of the approach used in the generation of promoters. **b.** Evaluation of the expression strength of promoters generated by PromoGen2-bsu in *B. subtilis*. **c.** Comparative analysis of the expression activities between promoters generated by PromoGen2-bsu and PromoGen-bsu in *B. subtilis*. **d.** Assessment of the expression strength of promoters generated by PromoGen2-bsu in *E. coli*. **e.** Comparative analysis of the expression activities between promoters generated by PromoGen2-bsu and PromoGen-bsu in *E. coli*. **f.** Evaluation of the transcriptional strength of promoters generated by PromoGen2-bsu in *B. licheniformis*. **g.** The correlation analysis of expression intensity between *B. subtilis* and *E. coli*, as well as between *B. subtilis* and *B. licheniformis*. **h.** Predictions by AlphaFold3 on the binding interactions between RNA polymerase and the generated promoters A2 and A4.

We analyzed the activity of the same native promoter sequence in *B. subtilis*, *E. coli*, and *P. aeruginosa*, finding that promoters active in low GC content *B. subtilis* were more likely to be active in other species (**Supplementary** Fig. 9, 10, and **11**). Therefore, the *B. subtilis* promoters generated by PromoGen2-bsu might be active in both Gram-positive and Gram-negative bacteria, showing potential cross-species applicability. To confirm this, we validated the activity of the 50 sequences active in *B. subtilis* in model organisms *E. coli* and *B. licheniformis*. Results showed that 38 out of the 50 sequences have higher promoter activity than the control promoter J23119 (**Fig. 4d**), while 100% of these tested sequences exhibited activity in *E. coli* and the average activity is 2.59-fold as high as the one of PromoGen generated promoters (**Fig. 4e**). In *B. licheniformis*, 48 out of 50 sequences exhibited activity and eighteen generated promoters were stronger than the control promoter *P_43_* (**Fig. 4f**). Moreover, the activity of the same promoter shows a strong correlation across different species. On the other hand, the Spearman correlation coefficient for the expression intensity of the generated promoter in *B. subtilis* and *E. coli* is 0.53, while the coefficient between *B. subtilis* and *B. licheniformis* is 0.92. Additionally, the expression intensity of promoters is more similar among closely related species, particularly for low-activity promoters (**Fig. 4g**). These findings indicated that promoters generated by the PromoGen2 model, fine-tuned with low GC content *B. subtilis* data, could serve as universal promoters across multiple species. In addition, we used the AlphaFold 3 webserver^32^ to predict the complex of PromoGen2 designed promoter and *E. coli* RNAP, the complex is structurally identical with X-ray solved structure (PDB ID: 4ynl). The predicted structure indicated that the highly active A4 promoter forms a stable RNAP-promoter open complex, where the −10 region is unwound within RNAP, facilitating transcription initiation. In contrast, the low-activity A2 promoter binds to RNAP in an inappropriate position, leaving the −10 region in a double-helical state, which hinders transcription initiation (**Fig. 4h**).

### *De novo* promoter generation based on the genome of understudied species

The exploration of understudied microbial cell factories, e.g. osmotic-pressure-tolerant gut bacterium *Jejubacter sp*. L23^33^, in metabolic engineering is limited by the lack of known regulatory elements to achieve suitable expression control, which could be solved by Promoter-Factory (**Fig. 5a**) in the absence of requiring any promoter-related experimental data. From the genome of the L23 strain, we used the Promoter-Factory to generate two models, differing in whether the base model was pre-trained, named PromoGen2-L23 and TM-L23, respectively. PromoGen2-L23 generated sequences showed high similarity as native promoters by 6-mer frequency analysis with Spearman and Pearson correlations of 0.764 and 0.881, respectively (**Fig. 5b**), outperforming the TM-L23 with correlations of 0.640 and 0.803, respectively. Assessed by PWM, the distribution of motifs generated by PromoGen2-L23 was highly consistent with the native sequences, showing a distinct TAATAT feature in the −10 region and no significant preferences in other regions (**Supplementary Fig. 12**) and indicating that PromoGen2-L23 successfully captured the characteristics and distribution patterns of L23 native promoters.

**Fig. 5.**
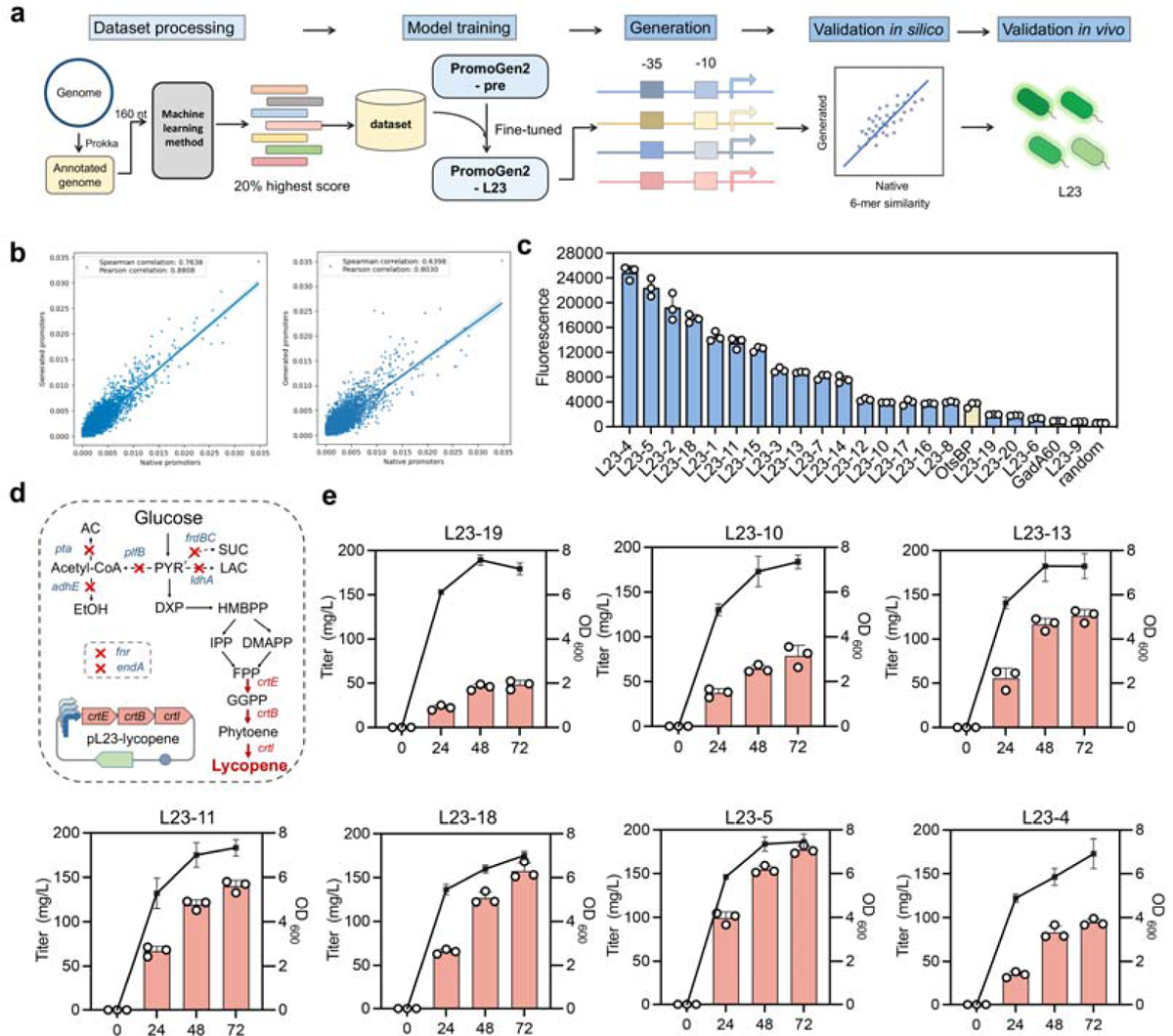
Training and validation of models for promoter generation in newly isolated bacterium *Jejubacter sp*. L23. **a.** Detailed description of the process in model training and validation. **b.** Spearman’s and Pearson’s r were utilized to assess the capability between the frequencies of 6-mers in native and generated promoters within L23. **c.** The expression strength of promoters generated by PromoGen2-L23 was evaluated within L23. **d.** The metabolic pathway of lycopene in L23R7. Seven genes were knockout including *frdBC* (fumarate reductase), *adhE* (alcohol dehydrogenase), *fnr* (fumarate and nitrate reductase), *pta* (phosphate acetyltransferase), *ldhA* (lactate dehydrogenase), *pflB* (pyruvate formate-lyase), and *endA* (Endonuclease). Three genes from *Pantoea ananatis* were expressed in L23R7 including geranylgeranyl pyrophosphate (GGPP) synthase, phytoene synthase, and phytoene desaturase. **e.** The production of lycopene in the L23R7 strain using promoters generated by PromoGen2.

Experimental results, using a random sequence as a negative control, showed that 95% and 80% of PromoGen2-L23-generated sequences (**Supplementary Table 4**) had higher expression than positive controls *P_gadA60_* and *P_otsBP_*, respectively, with regulatory ranges up to 37.4-fold and activity up to 25.7-fold as high as the one of *P_gadA60_* (**Fig. 5c**). BLAST analysis confirmed that no similarity existed between the PromoGen2-L23 generated promoters and the L23 genome, indicating that the sequences were *de novo* generated by the model based on learned promoter characteristics of L23, rather than resulting from data leakage.

The PromoGen2-L23 generated promoters were employed to drive the microbial production of linear C40 carotenoid lycopene (**Fig. 5d**), which is widely used as an antioxidant in functional foods, cosmetics, and pharmaceuticals, in L23R7, a genetically modified L23 variant. After 72 hours of shake-flask fermentation, the yields with L23-19, L23-10, L23-13, L23-11, L23-18, L23-5, and L23-4 promoters were 38.1, 78.8, 126.2, 140.1, 157.5, 177.1, and 94.4 mg/L, respectively (**Fig. 5e**), showing a 3.68-fold yield range.

### Training PromoGen2-proka for conditional design of promoters across thousands of prokaryotic genera

Based on the successful application of Promoter-Factory on L23, we aimed to further extend the applicability of customizable promoter generation to a broader range of species. To this end, a conditional generative model was developed to avoid the lab intensive customized fine-tuning process for individual species (**Fig. 6a**). To achieve this goal, we first constructed a cross-species core promoter dataset covering 5,800 prokaryotic genomes and 1,757 genera, for a total of 427,477 core promoter sequences, achieving extensive coverage of prokaryotic species (**Fig. 6b**), which was annotated through a combination of deep learning and interpretable machine learning methods. Based on this dataset, we conditionally fine-tuned the PromoGen2-base model, resulting in the PromoGen2-proka model, which can generate promoter sequences following a given genus prompt. We generated 1200 sequences for each of the 1,757 genera and created corresponding sequence logos, revealing variations in conserved features among different prokaryotic genera (**Supplementary Fig. 13**). Analysis of the Spearman’s r and Pearson’s r distributions between 4-mer frequencies in generated sequences and training data (**Fig. 6c**) showed that PromoGen2-proka effectively follows genus prompts, producing sequences highly consistent with native sequences from the target species. Further analysis of GC content differences and promoter sequence similarity between bacteria and archaea revealed that the 4-mer frequency similarity in the generated promoters was negatively correlated with the species differences of the GC content (**Fig. 6d**). This suggested that promoter prediction methods based on 4-mer frequency may exhibit limited generalizability across species with significant GC content divergence from the training data. Compared to phylogenetic distances based on 16S RNA sequences, the correlation between genome GC content differences and promoter sequence divergence was stronger (**Supplementary Fig. 14**), potentially indicating a co-evolutionary relationship between genomic GC content and promoter sequence patterns. Using *E. coli* as an example, we prompted PromoGen2-proka to generate sequences, which were subsequently validated experimentally, demonstrating transcriptional activities in all cases (**Fig. 6e**). These results indicated that genus-based prompting can be used for prokaryotic promoter generation.

**Fig. 6.**
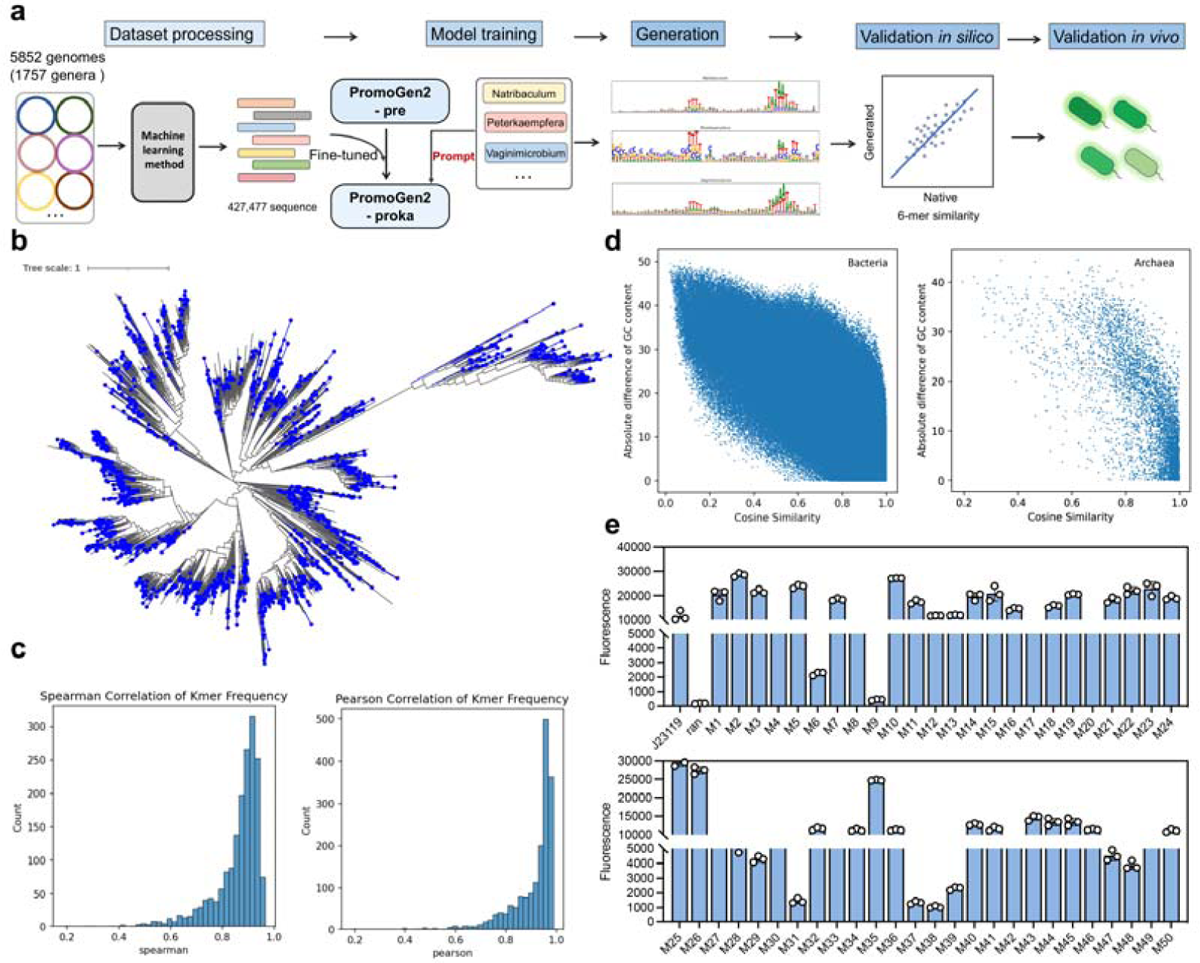
Training and validation of the PromoGen2-proka in various genera. **a.** Detailed description of the process in PromoGen2-proka training and validation. **b.** PromoGen2-proka spans the evolutionary tree of prokaryotes, with blue dots indicating the species we covered. **c.** Spearman and Pearson correlation distributions of k-mer frequencies across 1,757 genera. **d.** Relationship between k-mer similarity and species GC content. **e.** Validation of the activity of PromoGen2-proka-generated promoters in *E. coli*.

## Discussion

Various language models have successfully generated functional biomolecules, such as lysozymes^29^, P450 enzymes^34^, ribozymes^35^, and CREs like enhancers^36^ and promoters^23^. However, unlike protein and RNA sequences where reliable sequence-based structural and functional classification methods ensured vast growing training data^37^, the discoveries of CREs were heavily relied on experimental characterization. To address the challenge of design CREs in understudied species, we developed the Promoter-Factory, a computational protocol based on the PromoGen2 language model. PromoGen2 is the primary driver of Promoter-Factory’s success, with its effectiveness stemming from our meticulously curated training data. Furthermore, we suggest that zero-shot promoter strength prediction can serve as a benchmark for evaluating the suitability of generative language models as foundational models for CRE design. The generated promoters were successfully integrated into metabolic engineering efforts in a halophilic bacterium L23 for proof-of-concept lycopene production, extending the application of generated promoter elements to metabolic engineering for high-value compound production.

There is a clear demand for customized CREs with specific regulatory properties or desired expression strengths. Given the strong zero-shot generation capabilities of current language models—capable of accurately generating downstream sequences based on diverse prompts—and the architectural parallels between nucleotide language models and large-scale language models, we anticipate that with the integration of species-specific information and experimentally validated activity data, nucleic acid language models will achieve zero-shot generation of CREs with tailored strengths for individual species. Our development of a broadly applicable promoter design strategy represents a promising starting point for integrating generative-based promoter engineering into real-world synthetic biology applications.

While this work presents a library of promoters with varying strengths for a given species, the final target would be the generation of promoter sequences tailored to specific expression level within a given strain. However, this remains a challenge due to limited experimental data and the high costs associated with synthesizing large numbers of generated sequences. The future adaptation of PromoGen2 with manufacturing-aware generative models holds potential for enabling automatic petabase-scale synthesis of controllable promoters via an unbiased and synthesized oligo pool in the absence of cost-intensive experimental data^38^. Furthermore, such could expedite the identification of functional promoters across diverse species, reducing both the time and cost of discovery. This would also mitigate the limitations in promoter design quality associated with training on synthetic strain-specific data.

Taken together, we employed machine learning models to analyze microbial genomic data and constructed a DNA sequence dataset enriched with high-strength promoter sequences. Utilizing this dataset, we developed a nucleotide language model, PromoGen2, to decode prokaryotic CREs. PromoGen2 exhibited exceptional zero-shot promoter strength prediction accuracy, validated across both model and non-model organisms, underscoring the broad applicability of our approach to prokaryotic CRE design. We demonstrated that it is possible to train a generative model for a certain type of CREs even when experimental dataset is lack and the CREs could not be annotated in a homology-based way with model selected data. This approach showed great potential for accelerating the design of other CREs in the future.

## Methods

### Datasets for training

The prokaryotic genomes used for data collection were downloaded from NCBI, with only annotated genomes in RefSeq with assemblies as contigs or complete being selected to ensure quality. From the start position of each coding sequence (CDS) on the genome, an upstream sequence spanning 160 nucleotides was extracted. In total, 59 million sequences were gathered at this stage and served as the full collection of the prokaryotic untranslated region dataset (PURD). The data processing was performed using in-house Python scripts, utilizing the Biopython package. MMseqs2^39^ (v 13.45111) was used to remove redundancy from these sequences, with parameters of easy-cluster set to --min-id 0.9 and -c 0.9, resulting in 16 million clusters served as the PURD-90. For the representative sequences of each cluster, a previously trained CNN model for predicting transcriptional strength was applied, and sequences with predicted transcriptional strength greater than 2 were retained as a core collection of PURD, termed PURD-core. A total of 1.4 million sequences were kept in the PURD-core and used as pre-training data for the PromoGen2.

For fine-tuning the data used to train PromoGen2-PPD, the training data was downloaded from the PPD website. Sequences with identified TSSs and longer than 60 bp were retained for each organism. The genome files for the strains included in the PPD database were downloaded from NCBI, and all genomes were re-annotated using Prokka^40^ (v 1.14.6) to be aligned with the steps used in Promoter-Factory. The extraction of the core promoter regions was performed using the Promoter Calculator^13^.

### Generative language modeling

The PromoGen2 is a generative pretrained transformer model. During the pretraining stage of PromoGen2, including the PromoGen2-xsmall, PromoGen2-small, and the PromoGen2-base, the goal is to capture *P*(*b*), which *b* = (b_1_, …, b_n_) be a sequence of nucleotides and capped with the start token, five-prime and tailed with a three-prime token and the end of sequence token. Generative pretraining transformed sampling *b* from *P*(*b*) into the next-token prediction. The model with parameters θ was trained to minimize the negative log-likelihood over the training dataset

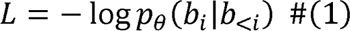

A new sample *b* of length *n* can be generated by autoregressive sampling from a five-prime token. The generation is expected to be stopped when an end of sequence token was sampled. The no-tag series models fine-tuned on each species followed the setting of pre-training stage. For conditional generative models including the tagged series models fine-tuned on each species and the multi-species models PromoGen2-xs-PPD, PromoGen2-s-PPD, and PromoGen2-b-PPD, a set of control tags *s* = (s_1_, …, s_m_) were used to represent each species used. Here, *x* = [*s*; *b*] be sequences inserted with a control tag between the start of sequence token and the five-prime token. The fine-tuning is to modify the pretrained parameters θ and find a new set of parameters θ’ to minimize the negative log-likelihood over the new dataset of *x*:

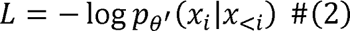

With θ’, a new promoter sequence *x* of length *n* can be sampled given the desired species encoded by the control tag *s* autoregressively: *pθ^’^(x_1_|s^-^), pθ^’^(x_2_|x^^^_1_, s^-^), …, pθ^’^(x_j_|x^^^_j_, s^-^)*

We used the decoder-only Transformer architecture for PromoGen2. In a Transformer model, multiple identical blocks are stacked to progressively update the representation at each position in the sequence. In each block, every position in the sequence is mapped to a hidden representation, and the relationships between these positions are inferred and updated accordingly using the self-attention mechanism.

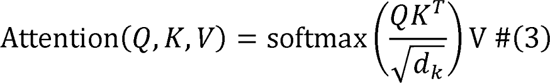

We applied multi-head self-attention to allow the neural network to capture different aspects of relationships between each position.

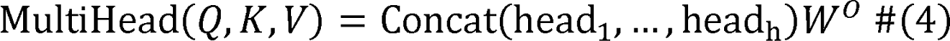

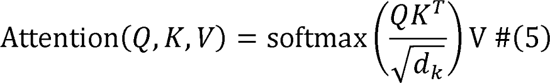

Since the self-attention mechanism itself lacks any notion of positional order, we adopted the positional encoding method used in GPT-2. Each position *i* in the sequence is assigned a unique positional embedding *E_pos_*(*i*) which is added to the word embedding:

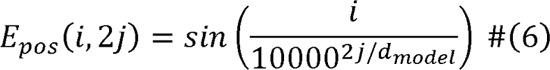

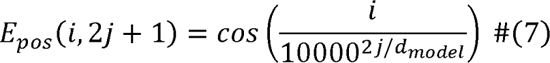

where j is the index of dimension, d_model_ is the hidden size.

The model was implemented with the transformers (v 4.38.2) package. The PromoGen2-xsmall, PromoGen2-small, and PromoGen2-base has 6, 12 and 30 blocks and the dimension of word embeddings is 320, 480, and 640 respectively.

### Training of the pretrained PromoGen2

During the pretraining, all models were trained for three epochs. Training was performed on a single RTX A6000 GPU and took for 2 days for the PromoGen2-base. Perplexity for the pre-trained models were evaluated on a holdout test set. The test perplexity did not exceed the training perplexity. The batch size was 64 sequences and all input sequences were truncated or padded to the length of 256. The Adam optimizer was used with learning rate of 1e^-4^. The learning rate was scheduled linearly to increase from 0 to 1e^-4^ over 50,00 initial steps and linear decay during the rest of the training. For the fine-tuning, we reduced the learning rate to 1e-5 and batch size to 32, with a reduced warmup step to 100 or the number of steps required by the first epoch, which is larger. The checkpoint with lowest validation loss were taken as the final weights.

### Implementation of the Promoter-Factory

For the generation of promoter sequences of a specific species beyond those included in the PPD, the pretrained PromoGen2 would require specific fine-tuning. The Promoter-Factory is designed to make this process more easily accessible without the need to write code. We decomposed the process to three key steps: 1) preparation of fine-tuning dataset, 2) model fine-tuning, 3) generation and evaluation. For the preparation of fine-tuning dataset, we used the prokka to annotate the CDS of the given genome. Using the annotated genbank file, the 160bp upstream sequence of each CDS were extracted. All sequences were scanned with a window size of 60 bp using Promoter Calculator to predict the transcript rate (Tx) with the last position as TSS. The window predicted with the highest Tx were kept as the core promoter. Core promoter ranked top 25% by the predicted Tx were served as the fine-tuning dataset. For model fine-tuning, the default setting is to prepend a user defined species token for conditional generative modeling. Other hyper-parameters were initialized identical to the fine-tuning procedure described in the neural network training section. Once the fine-tuning is finished, user would be able to generate sequences with desired sampling strategy with adjustable top-p and temperature values from the GUI. The program would report the correlation of k-mer frequency between the generated samples and the training samples. The GUI is implemented with the streamlit (v 1.31.0) package, metrics were calculated with the scipy (v 1.12.0) package.

### The PromoGen2-proka training

A total of 5,505 bacterial and 347 archaeal genomes marked as “reference” and “complete” were downloaded from NCBI (data as of 2024-10-5). Core promoters were annotated following the method used in the Promoter-Factory, and each full-length upstream sequence containing a core promoter was further scored using a CNN model. For selecting the training dataset, all promoter sequences were aggregated by genus. Within each genus group, a subset representing the top 90th percentile CNN scores were chosen, with the top 256 sequences by promoter calculator score included. In total, 1,757 genera contributed to the dataset, resulting in 427,477 core promoter sequences comprising the PCP (prokaryotic core promoter, PCP) dataset, used to train the final PromoGen2-proka model.

PromoGen2-base was fine-tuned on the PCP dataset, with 20,000 samples randomly selected as the test dataset. The learning rate was set to 1×10^-4^, weight decay was set to 0.1, and batch size was set to 128, with a warm-up period of 1,000 steps. The checkpoint with the lowest loss on the test dataset was retained as the final model. Training was conducted on an A100 GPU using fp16 precision.

### Sequence generation and analysis

Using the trained PromoGen2-proka, 1,200 promoter sequences were generated per genus. Sequences were filtered to retain only those with exactly 60 nucleotides. The prompt used for generating *E. coli* promoters for experimental testing was “<|bos|><|escherichia|>5.”

The sequence logos of generated sequences were analyzed using the WebLogo^41^ (v 3.7.12) package. For prokaryotic phylogenetic tree construction, 16S sequences from the NCBI 16S RefSeq records processing and curation were used. One representative 16S sequence per genus was randomly selected. Multiple sequence alignments were conducted using MAFFT^42^ (v7.526), and phylogenetic trees were constructed with FastTree^43^ (v 2.1.10) and visualized in iTol^44^. The GC content, homopolymer frequency analysis were performed with in-house python scripts. The CNN model used to predict promoter strength was developed in our previous study.

### Zero-shot prediction and metrics

We computed a log-likelihood score as the prediction of the strength of each sequence

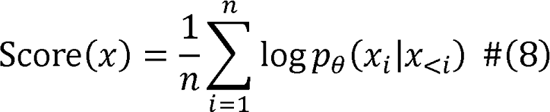

For BERT like model trained the masked language modeling objective, we compute the cross-entropy loss score instead of the log-likelihood score

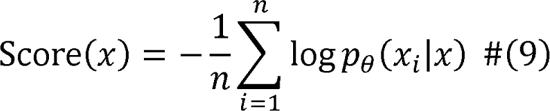

To measure the monotonic relationship between the predicted sequence strength and the ground truth, we used Spearman’s Rank Correlation Coefficient (ρ)

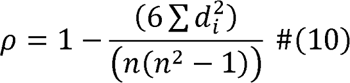

For measuring the linear relationship between the predicted sequence strength and the ground truth, we used Pearson’s Correlation Coefficient (r).

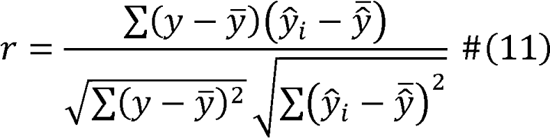

In the AUROC curve, the larger the area under the curve, the more accurate the classification. The x-axis represents the False Positive Rate (FPR), and the y-axis represents the True Positive Rate (TPR).

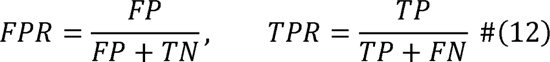

FP is the number of negative samples that are incorrectly classified as positive by the model. TN is the number of negative samples that are correctly classified as negative. TP is the number of positive samples that are correctly classified as positive. FN is the number of positive samples that are incorrectly classified as negative.

### Plasmid construction

The plasmids and primers used in this study are listed in Supplementary Table 5 and 6, respectively. All plasmid constructions were performed in *E. coli* JM109. The plasmids used in *B. subtilis* and *B. licheniformis* were based on the shuttle plasmid pXY43-sfGFP, which carries two origin of replication sites including pBR322 for construction in *E. coli* and repA for replication in *B. subtilis* and *B. licheniformis*. Using primers, we introduced 50 generated promoters into the pXY43-sfGFP plasmid, replacing the P43 promoter with PromoGen2-bsu-generated promoters via PCR. The PCR products were purified, treated with *Dpn1*, and transformed into *E. coli* JM109. The plasmids used for fluorescence intensity measurement in L23 were constructed using the same method. For lycopene fermentation, plasmid construction utilized the lab-preserved pDT315 plasmid as a template. The replaced promoter regions were incorporated into the pDT315 plasmid using primers and PCR. All constructed plasmids were verified through PCR and Sanger sequencing of the transformants to ensure successful construction.

### Assay of generated promoter strength

The constructed plasmids were transformed into the respective host strains, with *B. subtilis* using chemical transformation and *B. licheniformis* and L23 using electroporation. Single colonies from the plates were inoculated into LB medium containing kanamycin resistance and incubated overnight at 37°C with shaking at 220 rpm. Subsequently, the seed culture was diluted 1:100 into fresh LB medium with the appropriate antibiotic and incubated at 37°C with shaking at 220 rpm for 8 h. The culture was then diluted to an OD_600_ between 0.2 and 0.8, and 200 μL was transferred to a 96-well plate. A microplate reader (BioTek) was used to measure the OD_600_ and the sfGFP expression levels of the strains. Fluorescence intensity was measured with an excitation wavelength of 488 nm and an emission wavelength of 507 nm. The relative fluorescence intensity was calculated as sfGFP/OD_600_. All experiments were performed in triplicate.

### Lycopene fermentation in shake flasks

Seven successfully constructed lycopene fermentation plasmids were transformed into the L23R7 strain. Single colonies were selected from agar plates and cultured overnight in 5 mL of LB liquid medium containing Amp to prepare seed cultures. Subsequently, 1% (v/v) of the seed culture was inoculated into 50 mL of M9 medium supplemented with 40 g/L glucose as the carbon source. The fermentation was conducted at 30°C and 220 rpm for 72 h. Samples of 1.2 mL were collected every 24 h to measure OD_600_ and lycopene production.

### Lycopene extraction and detection

Lycopene accumulates intracellularly and is extracted from the cells using acetone. Frist, 1 mL of the fermentation broth is centrifuged at 12,000 rpm for 5 min to collect the cell pellet. The cell pellet is then resuspended in 2 mL of distilled water and centrifuged again to obtain a clean cell pellet. Subsequently, 1 mL of acetone is added to the cell pellet, and the mixture is homogenized before being incubated in a 55°C water bath until the cell pellet turns from red to white. The supernatant is collected, and its absorbance is measured at 474 nm. The lycopene concentration in the fermentation broth is determined using a standard curve for lycopene.

## Data availability

All data used to create training and evaluation datasets are freely available from public sources. Genomic sequence data are available at https://www.ncbi.nlm.nih.gov/datasets/genome/. 16S ResSeq data are available at https://www.ncbi.nlm.nih.gov/refseq/targetedloci/16S_process/. Promoter sequences with measured strength are from the ref.^12^.

## Code availability

The code of Promoter-Factory and the pre-trained weights will be available upon publication.

## Acknowledgements

This work was supported by the National Key R&D Program of China (2021YFC2100500), and the Natural Science Foundation of China (grant number 32370095). Part of the experiments was carried out in the Biological & Medical Engineering Core Facilities of the Beijing Institute of Technology.

## Author contributions

Y.-X.H., S.Y.G., Y.X., and J.Y.S conceptualized the project. Y.X., and J.Y.S designed, built, and implemented deep models in this study. Y.X., X.W.D., and Z.Y.L carried out the wet experiments. J.Y.S and W.Y.S carried out bioinformatic analysis. Y.-X.H., S.Y.G., Y.X., and J.Y.S analyzed the experiments data, and wrote the manuscript.

