## Supplemental information for "Language model generates *cis-*regulatory elements across prokaryotes"

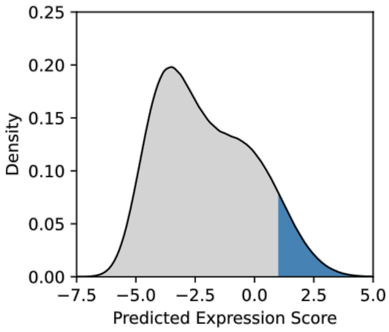


Figure S1. Scoring of core promoters by CNN.

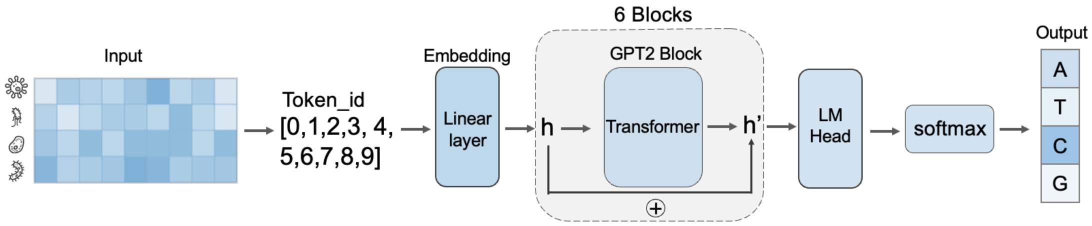


Figure S2. The model architecture of PromoGen2.

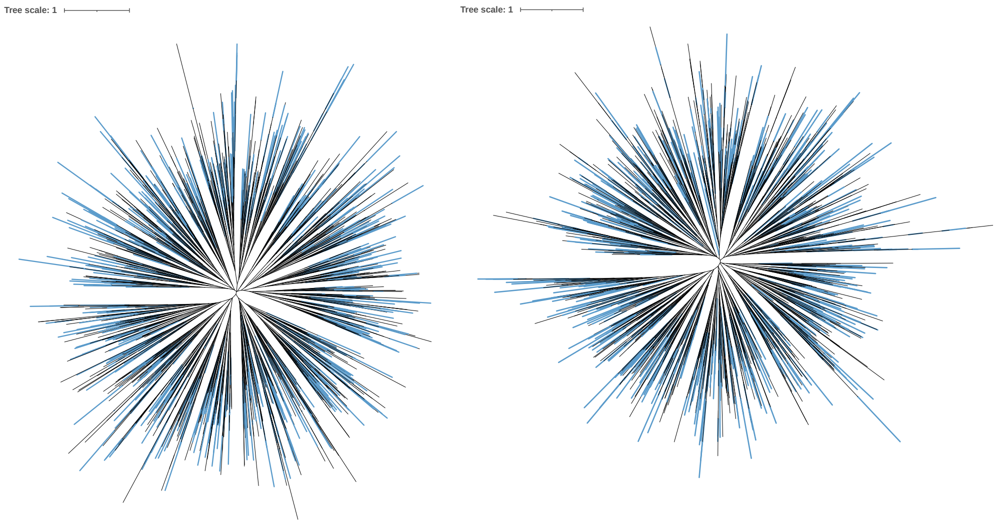


Figure S3. The Phylogenetic tree of PromoGen2-xsmall and -small generated sequence (blue) and native promoters (black).

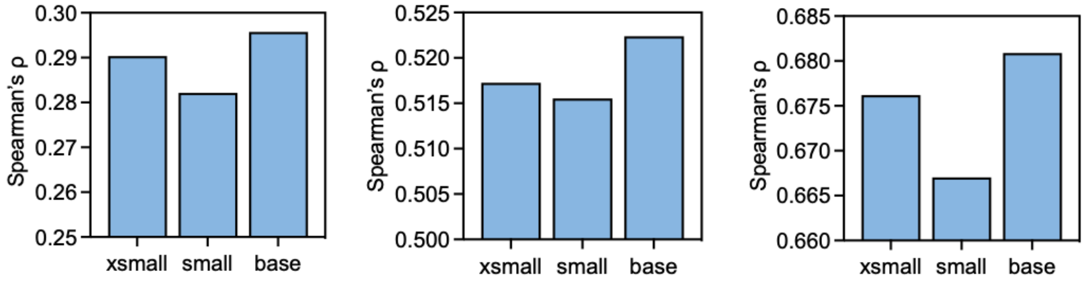


Figure S4. The Spearman correlation coefficients for *B. subtilis*, *E. coli and P. aeruginosa* in transcriptional levels.

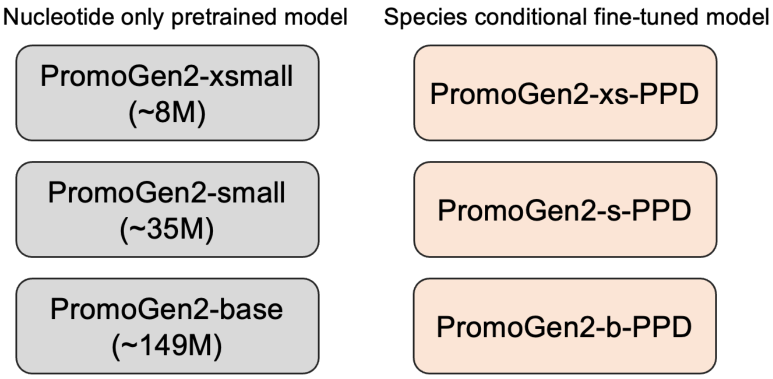


Figure S5. The finetuning models of PPD.

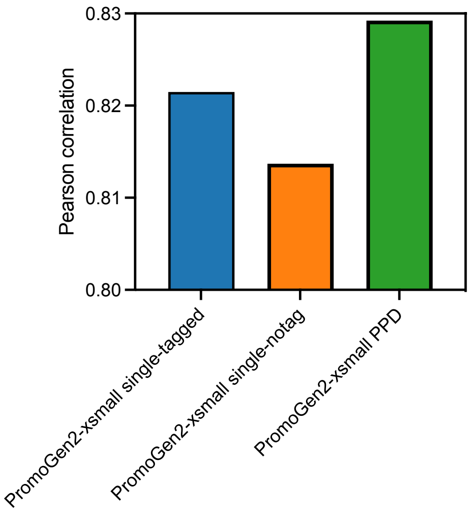


Figure S6. The average Pearson's correlation of the three finetuning models.

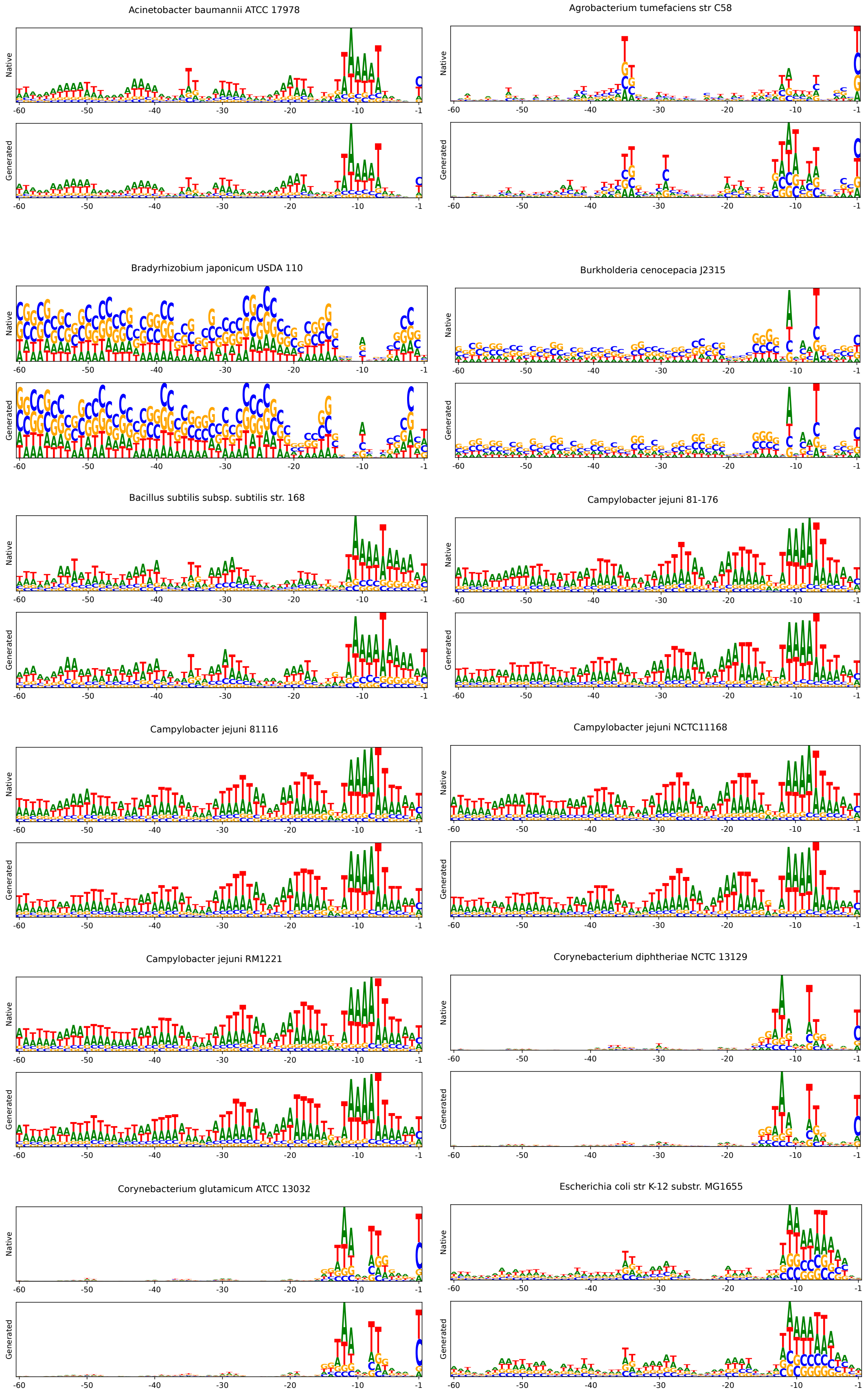
Acinetobacter baumannii ATCC 17978
Agrobacterium tumefaciens str C58
Bradyrhizobium japonicum USDA 110
Burkholderia cenocepacia J2315
Bacillus subtilis subsp. subtilis str. 168
Campylobacter jejuni 81-176
Campylobacter jejuni 81116
Campylobacter jejuni NCTC11168
Campylobacter jejuni RM1221
Corynebacterium diphtheriae NCTC 13129
Corynebacterium glutamicum ATCC 13032
Escherichia coli str K-12 substr. MG1655
Native
Generated
-60
-50
-40
-30
-20
-10
-1

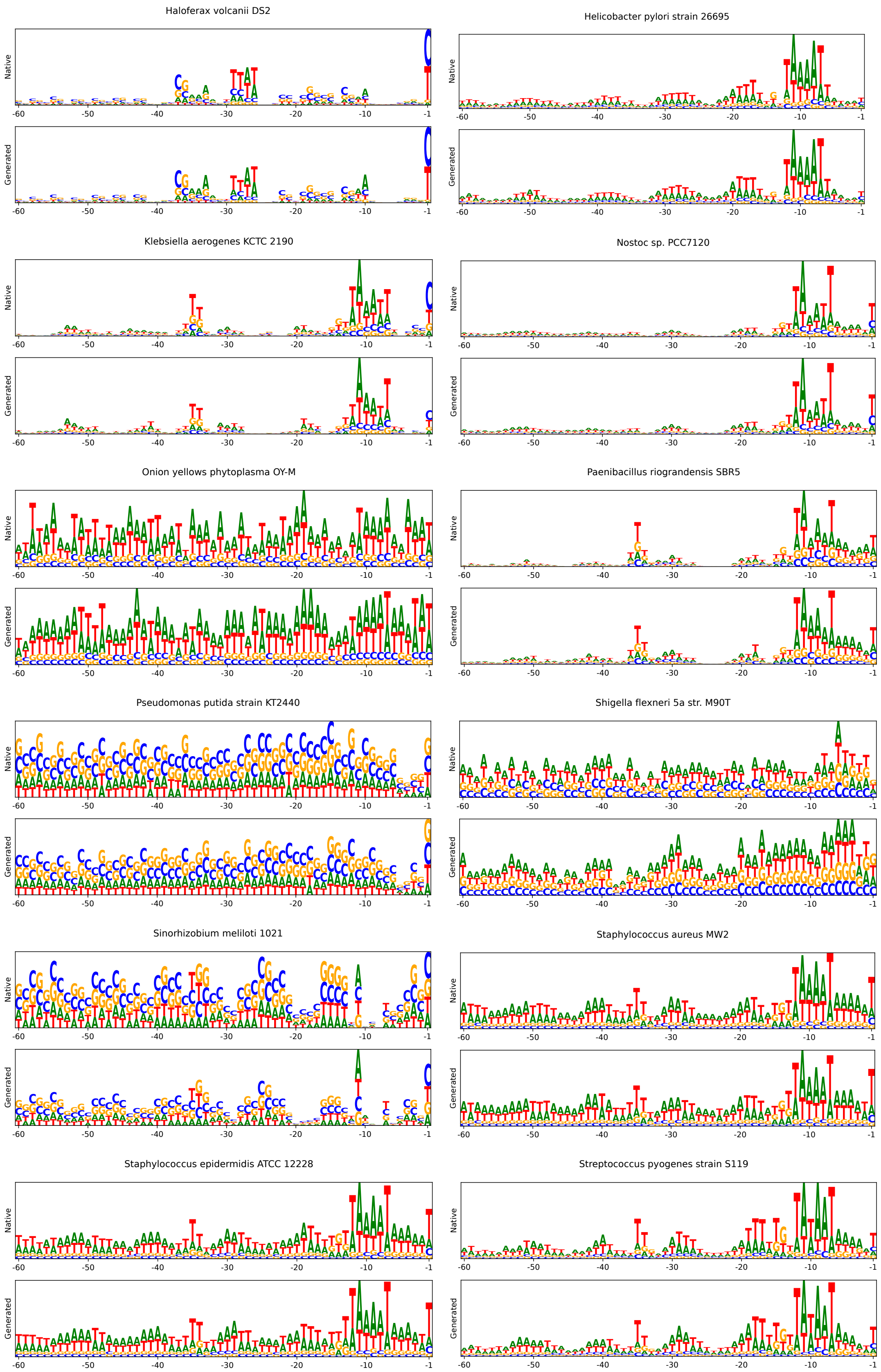
Haloferax volcanii DS2
Helicobacter pylori strain 26695
Klebsiella aerogenes KCTC 2190
Nostoc sp. PCC7120
Onion yellows phytoplasma OY-M
Paenibacillus riograndensis SBR5
Pseudomonas putida strain KT2440
Shigella flexneri 5a str. M90T
Sinorhizobium meliloti 1021
Staphylococcus aureus MW2
Staphylococcus epidermidis ATCC 12228
Streptococcus pyogenes strain S119
Native
Generated
-60
-50
-40
-30
-20
-10
-1

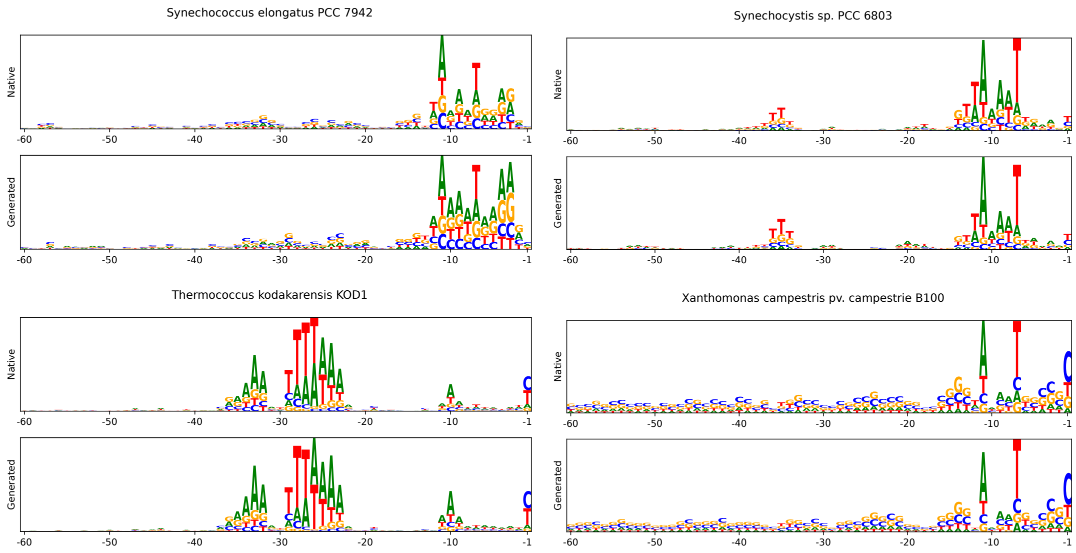


Figure S7. The sequence logos of the native, and PromoGen2-PPD-generated promoters.

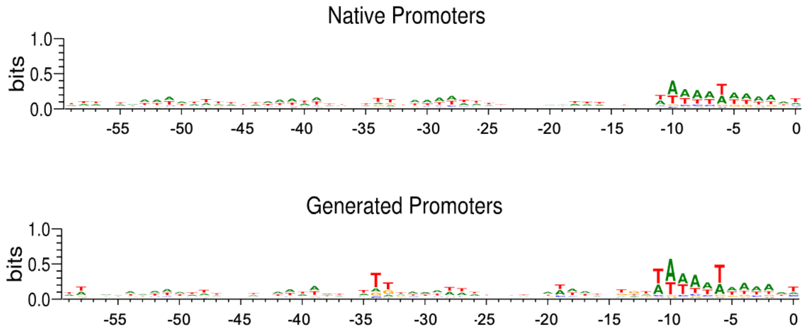


Figure S8. The sequence logos of the native, and PromoGen2-bsu-generated promoters.

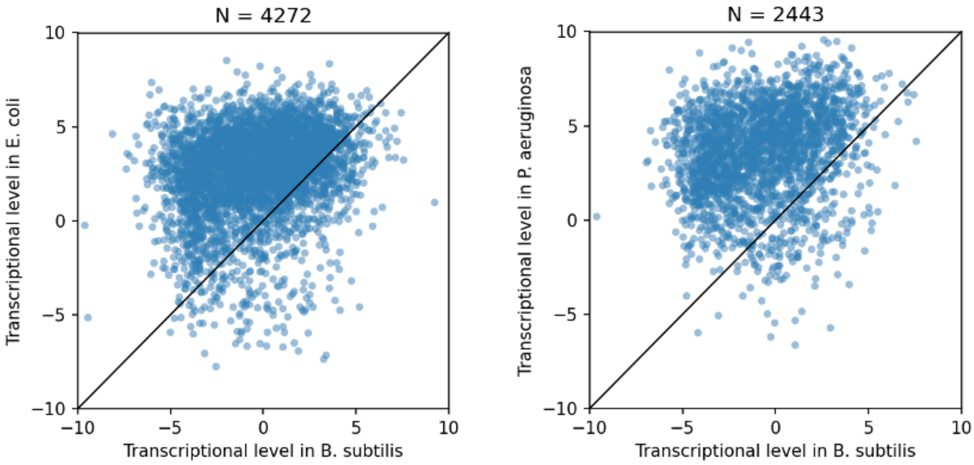

Figure S9. Analysis of the transcriptional activity of the same promoter sequences in *B. subtilis*, *E. coli* and *P. aeruginosa*.

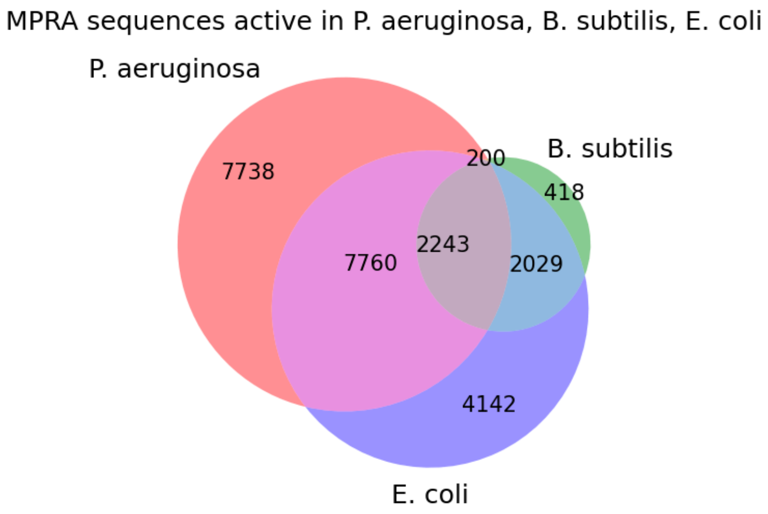


Figure S10. The promoter activities distribution in *E. coli*, *P. aeruginosa*, and *B. subtilis*.

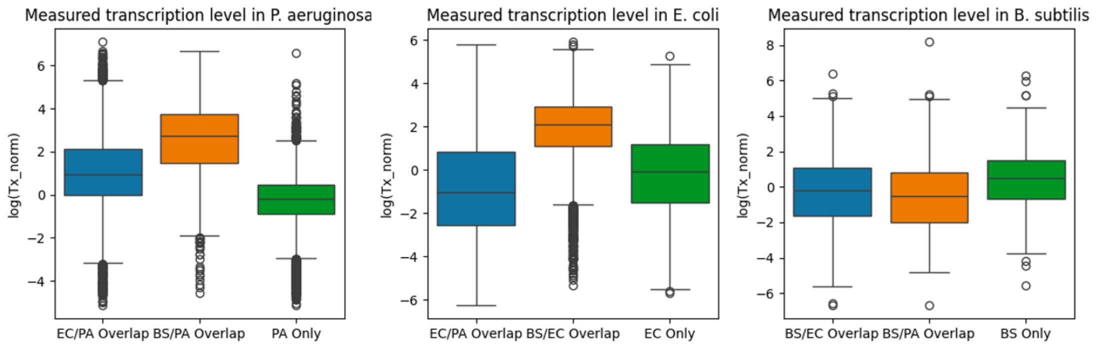


Figure S11. Analysis of the transcriptional activity of the same promoter sequences in *B. subtilis, E. coli* and *P. aeruginosa*.

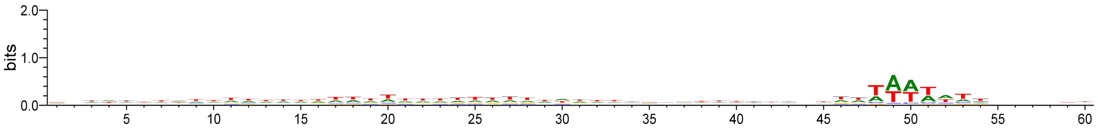


Figure S12. The sequence logos of the PromoGen2-L23-generated promoters.

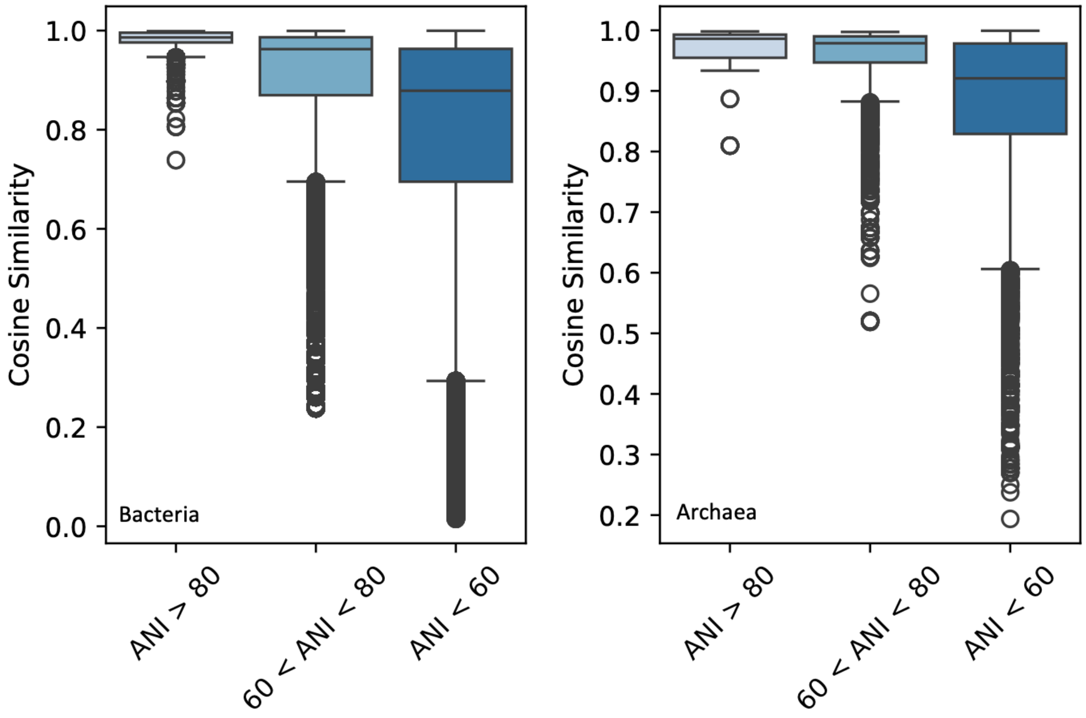


Figure S14. The relationship between evolutionary distance and promoter k-mer distribution among species.

**Table S1. The predictive performance scores of six models on promoter sequences.**

| | PA | SE | VN | EC | CG | BS |
|---|---|---|---|---|---|---|
| PG2-XS | 0.25549256 | 0.50342919 | 0.47236963 | 0.49404119 | 0.6157412 | 0.5113658 |
| PG2-S | 0.2464773 | 0.49676631 | 0.48048379 | 0.48205886 | 0.60889378 | 0.51613622 |
| PG2-B | 0.26300986 | 0.50988636 | 0.4736018 | 0.49683864 | 0.6245391 | 0.51482583 |
| NT-50M | 0.00151091 | 0.02878077 | 0.04611716 | -0.0209253 | 0.00507047 | 0.16790329 |
| NT-100M | 0.06997379 | 0.1119531 | 0.17147402 | 0.10162892 | 0.08027899 | -0.0071338 |
| DNABERT2 | -0.0104908 | 0.00144774 | 0.06673875 | 0.00214076 | 0.0571367 | -0.0200637 |

**Table S2. The Pearson correlation of the different models.**

| Model | PromoGen1 | PromoGen2-xsmall single-tagged | PromoGen2-xsmall single-notag | PromoGen2-xsmall PPD | Samples |
|---|---|---|---|---|---|
| *Campylobacter jejuni* NCTC11168 | 0.83735622 | 0.967640051 | 0.966857504 | 0.977850112 | 1905 |
| *Campylobacter jejuni* 81-176 | 0.85114941 | 0.967468985 | 0.959946626 | 0.974171104 | 2129 |
| *Campylobacter jejuni* RM1221 | 0.89023936 | 0.964802145 | 0.950829289 | 0.973406973 | 2166 |
| *Campylobacter jejuni* 81116 | 0.91197865 | 0.968002147 | 0.968884951 | 0.972130857 | 1942 |
| *Staphylococcus epidermidis* ATCC 12228 | 0.89105088 | 0.948890693 | 0.947352209 | 0.958533349 | 2207 |
| *Streptococcus pyogenes strain* S119 | 0.81742596 | 0.945473779 | 0.937625792 | 0.952685624 | 892 |
| *Staphylococcus aureus* MW2 | 0.89138208 | 0.951346183 | 0.952112588 | 0.952024525 | 2821 |
| *Helicobacter pylori* strain 26695 | 0.80399258 | 0.944831666 | 0.92987565 | 0.937386515 | 2233 |
| *Onion yellows phytoplasma* OY-M | 0.52619635 | 0.92578767 | 0.916292853 | 0.9187196 | 231 |
| *Bacillus subtilis* subsp. subtilis str. 168 | 0.7246909 | 0.920348816 | 0.917948616 | 0.905478063 | 691 |
| *Acinetobacter baumannii* ATCC 17978 | 0.78862513 | 0.909911531 | 0.900393321 | 0.90183415 | 1540 |
| *Burkholderia cenocepacia* J2315 | 0.78127671 | 0.861985974 | 0.892146108 | 0.891372733 | 10831 |
| *Thermococcus kodakarensis* KOD1 | 0.63325199 | 0.824783797 | 0.712851738 | 0.860936808 | 2720 |
| *Bradyrhizobium japonicum* USDA 110 | 0.69330331 | 0.831828959 | 0.839186626 | 0.853742571 | 15933 |
| *Xanthomonas campestris* pv. campestrie B100 | 0.70683558 | 0.836116804 | 0.842120601 | 0.846634597 | 3067 |

| | | | | | |
|---|---|---|---|---|---|
| *Nostoc sp.* PCC7120 | 0.64340336 | 0.820186913 | 0.849661746 | 0.835146742 | 13705 |
| *Pseudomonas putida* strain KT2440 | 0.63024428 | 0.839735501 | 0.824875645 | 0.83290325 | 7938 |
| *Sinorhizobium meliloti* 1021 | 0.64994342 | 0.804115334 | 0.80368975 | 0.828230368 | 17003 |
| *Haloferax volcanii* DS2 | 0.58636531 | 0.79732964 | 0.798739306 | 0.804132918 | 4749 |
| *Synechococcus elongatus* PCC 7942 | 0.44574692 | 0.624898455 | 0.531220245 | 0.781729703 | 1473 |
| *Escherichia coli* str K-12 substr. MG1655 | 0.53989724 | 0.710890913 | 0.75220864 | 0.775035245 | 8616 |
| *Synechocystis sp.* PCC 6803 | 0.42829189 | 0.766198216 | 0.764099456 | 0.763571644 | 944 |
| *Paenibacillus riograndensis* SBR5 | 0.4919874 | 0.753121023 | 0.731616339 | 0.735232759 | 2351 |
| *Shigella flexneri* 5a str. M90T | 0.47403116 | 0.717748573 | 0.756227035 | 0.705922425 | 14051 |
| *Klebsiella aerogenes* KCTC 2190 | 0.42555403 | 0.694276656 | 0.652312007 | 0.679813154 | 763 |
| *Agrobacterium tumefaciens* str C58 | 0.43499045 | 0.678973444 | 0.640989207 | 0.662218109 | 706 |
| *Corynebacterium diphtheriae* NCTC 13129 | 0.29039659 | 0.486066502 | 0.469591841 | 0.47041898 | 1656 |
| *Corynebacterium glutamicum* ATCC 13032 | 0.32071365 | 0.538917258 | 0.573710424 | 0.467156947 | 3581 |

**Table S3. The generated sequence by PromoGen2-bsu.**

| Name | Sequence (5'-3') |
|---|---|
| A1 | ATCTGAATTTTATTTATTTTAGATATTGATTTTAATTCACTATAGTGATATTATTAAAGT |
| A2 | AAATTATACAAGAAATTCAGTAAAATATAAATTTTACTGAATTTTATATATAATATAAAA |
| A3 | TTATTTGCTTTTTAATAATTATTGATTGAAATAAAAGAAAAATAGTGCTAATATATAGTT |
| A4 | AATGAAAGCGTTTTACTTAAATTATTTGAAATAATCTTTCAATTTGGTATAATAGATAAT |
| A5 | TGTTAAATTATAACATATTTGTGTATTGATAAACAATATGTTTATTGTTATAATGTATGT |
| A6 | TTGAACTTTAATTCAAAATATAATTTGAATTAGAGTTAAATTTATGGTAAAATGAATGCT |
| A7 | CATTATTGAATTTTTGTTCAATAATGTTTCAATAATATTGACTTATGATATAATTCTATT |
| A8 | TATAGGATTAATATATTGACAACAATATATTGATAAAATATAATGTATACATAAACAAAC |
| A9 | CTTTAGTTGAGTTTTTATTGATTATTAATTGAATATTAATTATTATAGTTATATAATATT |
| A10 | AGGAGGTTAAGTGGATAAAATTCGTTGACAACGAAAATGATTATCAATTACAATATAAAT |
| A11 | AATCACAAAAACTAAATATATATCCCTTGAATTATATTAAAACAAGCGTATAATACAGTT |
| A12 | AACCTTGATTTTTAGCTTGCTAAATATTGACAAAATTATTTTTAATTGTATAATTAGCTT |
| A13 | CGGAGATAAATATAGTATTTTTCTTTGTTTATATATTTAAAAATAGGTATAATAAAAGCT |
| A14 | TAATGAAAGACCAGTTAACGATATAGTTGATTGGTCTTTTTAATTTTGATAAGATAGATG |
| A15 | AAAAGAGAAGCCGTGCCCGGGCTTCTCTTTTATTTTGCCTTTCTCCGTTATAATGGCAAT |
| A16 | AATTATAAGGAAAAATACGGAACTTGTTTACATTTTAACAAAATAGGAATATAATGAAAT |
| A17 | AAGGAAAACGAATGGAAATCATTGCTTGACAAATTCATCATGTCCTGCTATACTTGTATA |
| A18 | TACACCATCTTATACATATCCTACTTTTTGTTTTATGTACATATATGATAAGATAATCTT |

| | |
|---|---|
| A19 | GTCAGAAAAAGTTAGCGTAAACAGTTTGCAATACACTACTTTTGGTGTAAAATCATAAGT |
| A20 | CGTCAGGTTTGGCGGTTTTTGTTTATTGATTTTCGGTCAGAATGGTGTTATACTAGATAC |
| A21 | ATGGGAGCGCAAAACACATTGATTTTTGCTTAGAACGAAGCTGAGTGGTATGCTTTATTG |
| A22 | TTTTACTAAAAATAATATTAACTGTTTTACAAAGTAATGCTAACTGTTATAATTTAAATT |
| A23 | TTCTTTATATTCTCTCTATTTTCTCTCATTTTCTCTCTATTTTCTCCTATTTGCCTCATT |
| A24 | TAGGACAAAGTTTAATTTATATAAAGTTGACAAAGATATAAAAAGTAGTATAATTATAAA |
| A25 | TTTTGGATAATAAATATCATAAATATTGATTATTAATTCAAATATAATTATACTATTTTT |
| A26 | TTTTAATATTAATTATTATGTTTATATACACTTTTAACAATAATGTGATATTATTGAATT |
| A27 | AATTTAACAAAATTATAAAATATGTAATTGAAAGCGCTATAAAATGATATAATAAGAATC |
| A28 | AAACCCCGAGAGGGGTTTTTGTTTTGGTTTGACTTTTTGATTAACTTTATATAAAATATA |
| A29 | TTGTAATTTTTATATAGAAAAATATTGAAATGAAAACCTTAAAGTGATATATTAAAAAGC |
| A30 | TTTGCTTTTAAAAATAAAATACTATTGACAATTATTATCAAATTAGAATAATCTAAATAT |
| A31 | TTCCATTTTTATTTTAAATTTGTCTATTCGAAAAAGAAAAATATGTGATATAATTATAAA |
| A32 | TATTTAAAAGCAAAATAAAAATAGATTGACAAATCGACTTTTTAGTAGTATACTTTCTTA |
| A33 | TATTTATATTTTATGATATGAAATGTCATATTTTATTTTTATTCTAGTAAAATATAAAAT |
| A34 | TATATCAAATTTTTCATAAAAAGCTTGACAAGTAGTGATTTATTGTTTATAGTATATATC |
| A35 | AGTGAAATATTTCACTAAAAATCTTGAAAACGCTTTAATGAATTCGGTATCATAGGTGTT |
| A36 | TTTAAAATTAGAATGTTATTTTTGTTGACAGATAAACCATTTGTTTATTATAATAAACAT |
| A37 | TAGTACTTTAAGCAACCGATTTAAATTGAAAAAGTCTATTTATAAGAGTAAAATGATATG |
| A38 | TCTTCACCTCGAGCACCTCTTTTCATTGTTAAGATTAGTATAGCATAGCTTTTCATAAAA |
| A39 | GTGAAGGATTTTACTAGAAAAAGACTTGTCAATATGAAAGTTTTCCTGTACTCTATAACT |
| A40 | TTCCACTCCTCTTAAAGAGGTCATACTTGACCTCAACTTTAAGTTTACTATACTTAATAT |
| A41 | TTGACAAAGCAATACCATCTGTATTATAATGTACAGATAGTATTTGTATATGAATATATA |
| A42 | AAAAGCCTACAAAGCAGCTGTTTTTGCCTTGACAACCGCCTAAGACAGTGTATACTAAGT |
| A43 | TATTCTTTTAAATACTCTTTTTACTTGAAAAATCAGCAGAGCTTATGTATAATTAATAAT |
| A44 | AATAGATATTTTAGCTTTATTTGTCCTTGCAAAAACCCAAAAACATGTTATACTTATCTT |
| A45 | TAATAGAATCATTCTATGGTTTTCTTATTGACTTTTAAAGCGGAATCGTTTACTATGATT |
| A46 | TTTCTAGAAAAGTTATTTACTTTTATTGCAAACTGTTTCGTCTTGTGATATTCTTTAATT |
| A47 | AAATAATAAAATCTTGATATATCGTTTGACAATAGAGTTATTAAGTGTTATTCTTGTAAC |
| A48 | TCCTCTAAATATTCTTTTTCTATTATTGATTTTTATTAAAAATAGGGGTACTATCATAAT |
| A49 | TTTTTAAAATCTAGGTATGAAAATAGCTTGAAATTCTGAAAATTAAGTGTTATATTTAAC |
| A50 | GTAAAATTTTGAATAAAAATTAGAATACTTTTGTTATGTAGTGTGATATAATAAGTAGTT |

**Table S4. The generated sequence by PromoGen2-L23**

| Name | Sequence (5'-3') |
|---|---|
| L23-1 | AAATAACATCACACACAAGCCGCCGATTGACAAAAGCGTGGCTTTTATATAGAATGCAGC |
| L23-2 | GATGCTTTCATTATTCTTTTTACTTCTTGATTTCCTCATACTGTTCAATATAATGAATTC |
| L23-3 | CAATTCAATTTGAATTGTCACTTGCATAAAAGTAGTGGTATTCATATTATTCAGTAAGTA |
| L23-4 | TGGCATACAAGCGCAAATATTCACTTGATTTATGCCAAAGAATGATGTATTATAATTACA |
| L23-5 | GACGACCACTGCCTGAATTTTTGGTTGAAATTGTCAATATCGCTTTGTAAAATCTGCGCG |
| L23-6 | ACTATTTTTACTATTTTAGCAATTTTAATTTGATAATTCATTGTGGTATTGCCTATATTT |
| L23-7 | TGCATCATTTTTGTGATGTTGCGATGTTGATATAAAAATCAACATCGTTATATTTTTGCG |
| L23-8 | GTAATATAATCATATTTCTCCCTTTGAAGATATACCGATATGGACTATATTTGCAGCCGT |
| L23-9 | TGAAAATAATAAAGAGTGATATAATTGACCGGTTTTATCACTCTTATTATAAATCAGTCC |
| L23-10 | AAAGTTTATTTATTCACAGCGTAATTGATAAATACAATCATTTATTTATAATCTACGTTA |
| L23-11 | ACCAATTTAAATAACAATACTTGATCAATTTACTCTGCACAATATATAATTACGCAAAGA |
| L23-12 | TATTAATATTATTGTAGGTTATTGATAAGTGTAAGATAATTCACTATTATTAATACTGTC |
| L23-13 | CAATTCAATTTGAATTGTCACTTGCATAAAAGTAGTGGTATTCATATTATTCAGTAAGTA |

| | |
|---|---|
| L23-14 | ATGAATCCTTTTTCATGACGCATTGAAATTTAATCACGCCGGTCATATTATGTATTTATT |
| L23-15 | TCACAATTTATAAAGTATTACCGCCTTGATGTGTAAAATTAATTACGCTACTATGCGGTT |
| L23-16 | AAATTTCTATCTTTACAACATTTTAGCAACCGCTATATATCATACTGCTATAATACTGAT |
| L23-17 | ATTTTTATCCTTTACATAAGTGATTGATTGGCAATATATTAAATTATATCATACCCCTTT |
| L23-18 | ACAATGCCGGTCTTCCGGATTGACTGGTAATTTGTGAATTATTGTGTTATAGTCCCGCCA |
| L23-19 | AGCCGGATGAATTATGGAGTTTTTGACAAATCGGCTCTGCTGGGCCGTATAATTGCCCCT |
| L23-20 | TAAATCGCTATGAAGGATTAAATGCCTGTTGAAAATTTATATTAACATATAATATACTTG |

**Table S5. The plasmids used in this study**

| Plasmids | Description | Source |
|---|---|---|
| pXY-P43-sfGFP | A plasmid expressing sfGFP under the control of the P43 promoter | Laboratory stock |
| pXY-PlepA-sfGFP | A plasmid expressing sfGFP under the control of the PlepA promoter | This study |
| pXY-Pveg-sfGFP | A plasmid expressing sfGFP under the control of the Pveg promoter | This study |
| pXY-A1-sfGFP | A plasmid expressing sfGFP under the control of the A1 promoter | This study |
| pXY-A2-sfGFP | A plasmid expressing sfGFP under the control of the A2 promoter | This study |
| pXY-A3-sfGFP | A plasmid expressing sfGFP under the control of the A3 promoter | This study |
| pXY-A4-sfGFP | A plasmid expressing sfGFP under the control of the A4 promoter | This study |
| pXY-A5-sfGFP | A plasmid expressing sfGFP under the control of the A5 promoter | This study |
| pXY-A6-sfGFP | A plasmid expressing sfGFP under the control of the A6 promoter | This study |
| pXY-A7-sfGFP | A plasmid expressing sfGFP under the control of the A7 promoter | This study |
| pXY-A8-sfGFP | A plasmid expressing sfGFP under the control of the A8 promoter | This study |
| pXY-A9-sfGFP | A plasmid expressing sfGFP under the control of the A9 promoter | This study |
| pXY-A10-sfGFP | A plasmid expressing sfGFP under the control of the A10 promoter | This study |
| pXY-A11-sfGFP | A plasmid expressing sfGFP under the control of the A11 promoter | This study |
| pXY-A12-sfGFP | A plasmid expressing sfGFP under the control of the A12 promoter | This study |
| pXY-A13-sfGFP | A plasmid expressing sfGFP under the control of the A13 promoter | This study |
| pXY-A14-sfGFP | A plasmid expressing sfGFP under the control of the A14 promoter | This study |
| pXY-A15-sfGFP | A plasmid expressing sfGFP under the control of the A15 promoter | This study |
| pXY-A16-sfGFP | A plasmid expressing sfGFP under the control of the A16 promoter | This study |

| | | |
|---|---|---|
| pXY-A17-sfGFP | A plasmid expressing sfGFP under the control of the A17 promoter | This study |
| pXY-A18-sfGFP | A plasmid expressing sfGFP under the control of the A18 promoter | This study |
| pXY-A19-sfGFP | A plasmid expressing sfGFP under the control of the A19 promoter | This study |
| pXY-A20-sfGFP | A plasmid expressing sfGFP under the control of the A20 promoter | This study |
| pXY-A21-sfGFP | A plasmid expressing sfGFP under the control of the A21 promoter | This study |
| pXY-A22-sfGFP | A plasmid expressing sfGFP under the control of the A22 promoter | This study |
| pXY-A23-sfGFP | A plasmid expressing sfGFP under the control of the A23 promoter | This study |
| pXY-A24-sfGFP | A plasmid expressing sfGFP under the control of the A24 promoter | This study |
| pXY-A25-sfGFP | A plasmid expressing sfGFP under the control of the A25 promoter | This study |
| pXY-A26-sfGFP | A plasmid expressing sfGFP under the control of the A26 promoter | This study |
| pXY-A27-sfGFP | A plasmid expressing sfGFP under the control of the A27 promoter | This study |
| pXY-A28-sfGFP | A plasmid expressing sfGFP under the control of the A28 promoter | This study |
| pXY-A29-sfGFP | A plasmid expressing sfGFP under the control of the A29 promoter | This study |
| pXY-A30-sfGFP | A plasmid expressing sfGFP under the control of the A30 promoter | This study |
| pXY-A31-sfGFP | A plasmid expressing sfGFP under the control of the A31 promoter | This study |
| pXY-A32-sfGFP | A plasmid expressing sfGFP under the control of the A32 promoter | This study |
| pXY-A33-sfGFP | A plasmid expressing sfGFP under the control of the A33 promoter | This study |
| pXY-A34-sfGFP | A plasmid expressing sfGFP under the control of the A34 promoter | This study |
| pXY-A35-sfGFP | A plasmid expressing sfGFP under the control of the A35 promoter | This study |
| pXY-A36-sfGFP | A plasmid expressing sfGFP under the control of the A36 promoter | This study |
| pXY-A37-sfGFP | A plasmid expressing sfGFP under the control of the A37 promoter | This study |
| pXY-A38-sfGFP | A plasmid expressing sfGFP under the control of the A38 promoter | This study |
| pXY-A39-sfGFP | A plasmid expressing sfGFP under the control of the A39 promoter | This study |
| pXY-A40-sfGFP | A plasmid expressing sfGFP under the control of the A40 promoter | This study |
| pXY-A41-sfGFP | A plasmid expressing sfGFP under the control of the A41 promoter | This study |

| | | |
|---|---|---|
| pXY-A42-sfGFP | A plasmid expressing sfGFP under the control of the A42 promoter | This study |
| pXY-A43-sfGFP | A plasmid expressing sfGFP under the control of the A43 promoter | This study |
| pXY-A44-sfGFP | A plasmid expressing sfGFP under the control of the A44 promoter | This study |
| pXY-A45-sfGFP | A plasmid expressing sfGFP under the control of the A45 promoter | This study |
| pXY-A46-sfGFP | A plasmid expressing sfGFP under the control of the A46 promoter | This study |
| pXY-A47-sfGFP | A plasmid expressing sfGFP under the control of the A47 promoter | This study |
| pXY-A48-sfGFP | A plasmid expressing sfGFP under the control of the A48 promoter | This study |
| pXY-A49-sfGFP | A plasmid expressing sfGFP under the control of the A49 promoter | This study |
| pXY-A50-sfGFP | A plasmid expressing sfGFP under the control of the A50 promoter | This study |
| pXY-L23-1-sfGFP | A plasmid expressing sfGFP under the control of the L23-1 promoter in L23 | This study |
| pXY-L23-2-sfGFP | A plasmid expressing sfGFP under the control of the L23-2 promoter in L23 | This study |
| pXY-L23-3-sfGFP | A plasmid expressing sfGFP under the control of the L23-3 promoter in L23 | This study |
| pXY-L23-4-sfGFP | A plasmid expressing sfGFP under the control of the L23-4 promoter in L23 | This study |
| pXY-L23-5-sfGFP | A plasmid expressing sfGFP under the control of the L23-5 promoter in L23 | This study |
| pXY-L23-6-sfGFP | A plasmid expressing sfGFP under the control of the L23-6 promoter in L23 | This study |
| pXY-L23-7-sfGFP | A plasmid expressing sfGFP under the control of the L23-7 promoter in L23 | This study |
| pXY-L23-8-sfGFP | A plasmid expressing sfGFP under the control of the L23-8 promoter in L23 | This study |
| pXY-L23-9-sfGFP | A plasmid expressing sfGFP under the control of the L23-9 promoter in L23 | This study |
| pXY-L23-10-sfGFP | A plasmid expressing sfGFP under the control of the L23-10 promoter in L23 | This study |
| pXY-L23-11-sfGFP | A plasmid expressing sfGFP under the control of the L23-11 promoter in L23 | This study |
| pXY-L23-12-sfGFP | A plasmid expressing sfGFP under the control of the L23-12 promoter in L23 | This study |
| pXY-L23-13-sfGFP | A plasmid expressing sfGFP under the control of the L23-13 promoter in L23 | This study |
| pXY-L23-14-sfGFP | A plasmid expressing sfGFP under the control of the L23-14 promoter in L23 | This study |
| pXY-L23-15-sfGFP | A plasmid expressing sfGFP under the control of the L23-15 promoter in L23 | This study |
| pXY-L23-16-sfGFP | A plasmid expressing sfGFP under the control of the L23-16 promoter in L23 | This study |

| | | |
|---|---|---|
| pXY-L23-17-sfGFP | A plasmid expressing sfGFP under the control of the L23-17 promoter in L23 | This study |
| pXY-L23-18-sfGFP | A plasmid expressing sfGFP under the control of the L23-18 promoter in L23 | This study |
| pXY-L23-19-sfGFP | A plasmid expressing sfGFP under the control of the L23-19 promoter in L23 | This study |
| pXY-L23-20-sfGFP | A plasmid expressing sfGFP under the control of the L23-20 promoter in L23 | This study |
| pXY-L23-19-crtEBI | The lycopene fermentation plasmid under the expression of Promoter L23-19 | This study |
| pXY-L23-10-crtEBI | The lycopene fermentation plasmid under the expression of Promoter L23-10 | This study |
| pXY-L23-13-crtEBI | The lycopene fermentation plasmid under the expression of Promoter L23-13 | This study |
| pXY-L23-11-crtEBI | The lycopene fermentation plasmid under the expression of Promoter L23-11 | This study |
| pXY-L23-18-crtEBI | The lycopene fermentation plasmid under the expression of Promoter L23-18 | This study |
| pXY-L23-5-crtEBI | The lycopene fermentation plasmid under the expression of Promoter L23-5 | This study |
| pXY-L23-4-crtEBI | The lycopene fermentation plasmid under the expression of Promoter L23-4 | This study |

**Table S6. The Primers used in this study.**

| **Primers** | **Sequences (5'-3')** |
|---|---|
| A1-F | ATATTGATTTTAATTCACTATAGTGATATTATTAAAGTAAAGGAGGTGATAAAAATGGTTAGC |
| A1-R | TAGTGAATTAAAATCAATATCTAAAATAAATAAAATTCAGATCTAGTTACCCTCGAGATCTGG |
| A2-F | AAATATAAATTTTACTGAATTTTATATATAATATAAAAAAAGGAGGTGATAAAAATGGTTAGC |
| A2-R | ATTCAGTAAAATTTATATTTTACTGAATTTCTTGTATAATTTCTAGTTACCCTCGAGATCTGG |
| A3-F | TGATTGAAATAAAAGAAAAATAGTGCTAATATATAGTTAAAGGAGGTGATAAAAATGGTTAGC |
| A3-R | TTTTTCTTTTATTTCAATCAATAATTATTAAAAAGCAAATAACTAGTTACCCTCGAGATCTGG |
| A4-F | TATTTGAAATAATCTTTCAATTTGGTATAATAGATAATAAAGGAGGTGATAAAAATGGTTAGC |
| A4-R | TTGAAAGATTATTTCAAATAATTTAAGTAAAACGCTTTCATTCTAGTTACCCTCGAGATCTGG |
| A5-F | GTATTGATAAACAATATGTTTATTGTTATAATGTATGTAAAGGAGGTGATAAAAATGGTTAGC |
| A5-R | AACATATTGTTTATCAATACACAAATATGTTATAATTTAACACTAGTTACCCTCGAGATCTGG |
| A6-F | AATTTGAATTAGAGTTAAATTTATGGTAAAATGAATGCTAAAGGAGGTGATAAAAATGGTTAGC |
| A6-R | AATTTAACTCTAATTCAAATTATATTTTGAATTAAAGTTCAACTAGTTACCCTCGAGATCTGG |

| | |
|---|---|
| A7-F | TAATGTTTCAATAATATTGACTTATGATATAATTCTATTAAAGGAGGTGATAAAAATGGTTAGC |
| A7-R | GTCAATATTATTGAAACATTATTGAACAAAAATTCAATAATGCTAGTTACCCTCGAGATCTGG |
| A8-F | CAATATATTGATAAAATATAATGTATACATAAACAAACAAAGGAGGTGATAAAAATGGTTAGC |
| A8-R | TATATTTTATCAATATATTGTTGTCAATATATTAATCCTATACTAGTTACCCTCGAGATCTGG |
| A9-F | TTATTAATTGAATATTAATTATTATAGTTATATAATATTAAAGGAGGTGATAAAAATGGTTAGC |
| A9-R | TAATTAATATTCAATTAATAATCAATAAAAACTCAACTAAAGCTAGTTACCCTCGAGATCTGG |
| A10-F | CGTTGACAACGAAAATGATTATCAATTACAATATAAATAAAGGAGGTGATAAAAATGGTTAGC |
| A10-R | AATCATTTTCGTTGTCAACGAATTTTATCCACTTAACCTCCTCTAGTTACCCTCGAGATCTGG |
| A11-F | ATCCCTTGAATTATATTAAAACAAGCGTATAATACAGTTAAAGGAGGTGATAAAAATGGTTAGC |
| A11-R | TTTTAATATAATTCAAGGGATATATATTTAGTTTTTGTGATTCTAGTTACCCTCGAGATCTGG |
| A12-F | TAAATATTGACAAAATTATTTTTAATTGTATAATTAGCTTAAAGGAGGTGATAAAAATGGTTAGC |
| A12-R | AAAATAATTTTGTCAATATTTAGCAAGCTAAAAATCAAGGTTCTAGTTACCCTCGAGATCTGG |
| A13-F | CTTTGTTTATATATTTAAAAATAGGTATAATAAAAGCTAAAGGAGGTGATAAAAATGGTTAGC |
| A13-R | TTTTAAATATATAAACAAAGAAAAATACTATATTTATCTCCGCTAGTTACCCTCGAGATCTGG |
| A14-F | ATAGTTGATTGGTCTTTTTAATTTTGATAAGATAGATGAAAGGAGGTGATAAAAATGGTTAGC |
| A14-R | TAAAAAGACCAATCAACTATATCGTTAACTGGTCTTTCATTACTAGTTACCCTCGAGATCTGG |
| A15-F | TTCTCTTTTATTTTGCCTTTCTCCGTTATAATGGCAATAAAGGAGGTGATAAAAATGGTTAGC |
| A15-R | AAAGGCAAAATAAAAGAGAAGCCCGGGCACGGCTTCTCTTTTCTAGTTACCCTCGAGATCTGG |
| A16-F | CTTGTTTACATTTTAACAAAATAGGAATATAATGAAATAAAGGAGGTGATAAAAATGGTTAGC |
| A16-R | TATTTTGTTAAAATGTAAACAAGTTCCGTATTTTTCCTTATAATTCTAGTTACCCTCGAGATCTGG |
| A17-F | TGCTTGACAAATTCATCATGTCCTGCTATACTTGTATAAAAGGAGGTGATAAAAATGGTTAGC |
| A17-R | CATGATGAATTTGTCAAGCAATGATTTCCATTCGTTTTCCTTCTAGTTACCCTCGAGATCTGG |
| A18-F | ACTTTTTGTTTTATGTACATATATGATAAGATAATCTTAAAGGAGGTGATAAAAATGGTTAGC |
| A18-R | ATGTACATAAAACAAAAAGTAGGATATGTATAAGATGGTGTACTAGTTACCCTCGAGATCTGG |
| A19-F | AGTTTGCAATACACTACTTTTGGTGTAAAATCATAAGTAAAGGAGGTGATAAAAATGGTTAGC |

| | |
|---|---|
| A19-R | AAAGTAGTGTATTGCAAACTGTTTACGCTAACTTTTTCTGACCTAGTTACCCTCGAGATCTGG |
| A20-F | TTATTGATTTTCGGTCAGAATGGTGTTATACTAGATACAAAGGAGGTGATAAAAATGGTTAGC |
| A20-R | TTCTGACCGAAAATCAATAAACAAAAACCGCCAAACCTGACGCTAGTTACCCTCGAGATCTGG |
| A21-F | TTTTTGCTTAGAACGAAGCTGAGTGGTATGCTTTATTGAAAGGAGGTGATAAAAATGGTTAGC |
| A21-R | AGCTTCGTTCTAAGCAAAAATCAATGTGTTTTGCGCTCCCATCTAGTTACCCTCGAGATCTGG |
| A22-F | TGTTTTACAAAGTAATGCTAACTGTTATAATTTAAATTAAAGGAGGTGATAAAAATGGTTAGC |
| A22-R | TAGCATTACTTTGTAAAACAGTTAATATTATTTTTAGTAAAACTAGTTACCCTCGAGATCTGG |
| A23-F | CTCTCATTTTCTCTCTATTTTCTCCTATTTGCCTCATTAAAGGAGGTGATAAAAATGGTTAGC |
| A23-R | AAATAGAGAGAAAATGAGAGAAAATAGAGAGAATATAAAGAACTAGTTACCCTCGAGATCTGG |
| A24-F | ATAAAGTTGACAAAGATATAAAAAGTAGTATAATTATAAAAAAGGAGGTGATAAAAATGGTTAGC |
| A24-R | TTTATATCTTTGTCAACTTTATATAAATTAAACTTTGTCCTACTAGTTACCCTCGAGATCTGG |
| A25-F | ATATTGATTATTAATTCAAATATAATTATACTATTTTTAAAGGAGGTGATAAAAATGGTTAGC |
| A25-R | TTTGAATTAATAATCAATATTTATGATATTTATTATCCAAAACTAGTTACCCTCGAGATCTGG |
| A26-F | TTATATACACTTTTAACAATAATGTGATATTATTGAATTAAAGGAGGTGATAAAAATGGTTAGC |
| A26-R | TATTGTTAAAAGTGTATATAAACATAATAATTAATATTAAAACTAGTTACCCTCGAGATCTGG |
| A27-F | TGTAATTGAAAGCGCTATAAAATGATATAATAAGAATCAAAGGAGGTGATAAAAATGGTTAGC |
| A27-R | TTATAGCGCTTTCAATTACATATTTTATAATTTTGTTAAATTCTAGTTACCCTCGAGATCTGG |
| A28-F | TTTTGGTTTGACTTTTTGATTAACTTTATATAAAATATAAAAGGAGGTGATAAAAATGGTTAGC |
| A28-R | AATCAAAAAGTCAAACCAAAACAAAAACCCCTCTCGGGGTTTCTAGTTACCCTCGAGATCTGG |
| A29-F | TATTGAAATGAAAACCTTAAAGTGATATATTAAAAAGCAAAGGAGGTGATAAAAATGGTTAGC |
| A29-R | TTAAGGTTTTCATTTCAATATTTTTCTATATAAAAATTACAACTAGTTACCCTCGAGATCTGG |
| A30-F | TATTGACAATTATTATCAAATTAGAATAATCTAAATATAAAGGAGGTGATAAAAATGGTTAGC |
| A30-R | TTTGATAATAATTGTCAATAGTATTTTATTTTTAAAAGCAAACTAGTTACCCTCGAGATCTGG |
| A31-F | GTCTATTCGAAAAAGAAAAATATGTGATATAATTATAAAAAAGGAGGTGATAAAAATGGTTAGC |
| A31-R | ATTTTTCTTTTTCGAATAGACAAATTTAAAATAAAAATGGAACTAGTTACCCTCGAGATCTGG |

| | |
|---|---|
| A32-F | TAGATTGACAAATCGACTTTTTAGTAGTATACTTTCTTAAAAGGAGGTGATAAAAATG<br>GTTAGC |
| A32-R | AAAAGTCGATTTGTCAATCTATTTTTATTTTGCTTTTAAATACTAGTTACCCTCGAGA<br>TCTGG |
| A33-F | ATGTCATATTTTATTTTTATTCTAGTAAAATATAAAATAAAGGAGGTGATAAAAATGG<br>TTAGC |
| A33-R | ATAAAAATAAAATATGACATTTCATATCATAAAATATAAATACTAGTTACCCTCGAGA<br>TCTGG |
| A34-F | AGCTTGACAAGTAGTGATTTATTGTTTATAGTATATATCAAAGGAGGTGATAAAAATG<br>GTTAGC |
| A34-R | TAAATCACTACTTGTCAAGCTTTTTATGAAAAATTTGATATACTAGTTACCCTCGAGA<br>TCTGG |
| A35-F | CTTGAAAACGCTTTAATGAATTCGGTATCATAGGTGTTAAAGGAGGTGATAAAAATGG<br>TTAGC |
| A35-R | TTCATTAAAGCGTTTTCAAGATTTTTAGTGAAATATTTCACTCTAGTTACCCTCGAGA<br>TCTGG |
| A36-F | TGTTGACAGATAAACCATTTGTTTATTATAATAAACATAAAGGAGGTGATAAAAATGG<br>TTAGC |
| A36-R | AAATGGTTTATCTGTCAACAAAAATAACATTCTAATTTTAAACTAGTTACCCTCGAGA<br>TCTGG |
| A37-F | AAATTGAAAAAGTCTATTTATAAGAGTAAAATGATATGAAAGGAGGTGATAAAAATGG<br>TTAGC |
| A37-R | TAAATAGACTTTTTCAATTTAAATCGGTTGCTTAAAGTACTACTAGTTACCCTCGAGA<br>TCTGG |
| A38-F | AACTTGACAGCGTATCATTTAAGTGCTATAATTCTATAAAAGGAGGTGATAAAAATGG<br>TTAGC |
| A38-R | AAATGATACGCTGTCAAGTTTAAGTATAGTAAGTATTCAAATCTAGTTACCCTCGAGA<br>TCTGG |
| A39-F | GACTTGTCAATATGAAAGTTTTCCTGTACTCTATAACTAAAGGAGGTGATAAAAATGG<br>TTAGC |
| A39-R | AACTTTCATATTGACAAGTCTTTTTCTAGTAAAATCCTTCACCTAGTTACCCTCGAGA<br>TCTGG |
| A40-F | ATACTTGACCTCAACTTTAAGTTTACTATACTTAATATAAAGGAGGTGATAAAAATGG<br>TTAGC |
| A40-R | TTAAAGTTGAGGTCAAGTATGACCTCTTTAAGAGGAGTGGAACTAGTTACCCTCGAGA<br>TCTGG |
| A41-F | ATGTTGAATTTTAACATAATACCTATTATACTGAACATTAAAGGAGGTGATAAAAATG<br>GTTAGC |
| A41-R | TATTATGTTAAAATTCAACATAATATTGCGAATTTCTTCAGACTAGTTACCCTCGAGA<br>TCTGG |
| A42-F | TTTGCCTTGACAACCGCCTAAGACAGTGTATACTAAGTAAAGGAGGTGATAAAAATGG<br>TTAGC |
| A42-R | TAGGCGGTTGTCAAGGCAAAAACAGCTGCTTTGTAGGCTTTTCTAGTTACCCTCGAGA<br>TCTGG |
| A43-F | ACTTGAAAAATCAGCAGAGCTTATGTATAATTAATAATTAAAGGAGGTGATAAAAATG<br>GTTAGC |
| A43-R | GCTCTGCTGATTTTTCAAGTAAAAAGAGTATTTAAAAGAATACTAGTTACCCTCGAGA<br>TCTGG |
| A44-F | TGTCCTTGCAAAAACCCAAAAACATGTTATACTTATCTTAAAGGAGGTGATAAAAATG<br>GTTAGC |

| | |
|---|---|
| A44-R | TTTTGGGTTTTTGCAAGGACAAATAAAGCTAAAATATCTATTCTAGTTACCCTCGAGATCTGG |
| A45-F | TCTTATTGACTTTTAAAGCGGAATCGTTTACTATGATTAAAGGAGGTGATAAAAATGGTTAGC |
| A45-R | CGCTTTAAAAGTCAATAAGAAAACCATAGAATGATTCTATTACTAGTTACCCTCGAGATCTGG |
| A46-F | TTATTGCAAACTGTTTCGTCTTGTGATATTCTTTAATTAAAGGAGGTGATAAAAATGGTTAGC |
| A46-R | GACGAAACAGTTTGCAATAAAAGTAAATAACTTTTCTAGAAACTAGTTACCCTCGAGATCTGG |
| A47-F | TCGTTTGACAATAGAGTTATTAAGTGTTATTCTTGTAACAAAGGAGGTGATAAAAATGGTTAGC |
| A47-R | AATAACTCTATTGTCAAACGATATATCAAGATTTTATTATTTCTAGTTACCCTCGAGATCTGG |
| A48-F | ATAGCTTGAAATTCTGAAAATTAAGTGTTATATTTAACAAAGGAGGTGATAAAAATGGTTAGC |
| A48-R | TTTTCAGAATTTCAAGCTATTTTCATACCTAGATTTTAAAAACTAGTTACCCTCGAGATCTGG |
| A49-F | GAATACTTTTGTTATGTAGTGTGATATAATAAGTAGTTAAAGGAGGTGATAAAAATGGTTAGC |
| A49-R | ACTACATAACAAAAGTATTCTAATTTTTATTCAAAATTTTACCTAGTTACCCTCGAGATCTGG |
| A50-F | ATGCTTGACAATACATTGTTAGCATTATAAAATGCAATCAAAGGAGGTGATAAAAATGGTTAGC |
| A50-R | TAACAATGTATTGTCAAGCATACATTAAGCTCTTAGCAATGTCTAGTTACCCTCGAGATCTGG |
| L23-1F | CCGATTGACAAAAGCGTGGCTTTTATATAGAATGCAGCAAAGGAGGTGATAAAAATGGTTAGC |
| L23-1R | GCCACGCTTTTGTCAATCGGCGGCTTGTGTGTGATGTTATTTCTAGTTACCCTCGAGATCTGG |
| L23-2F | CTTCTTGATTTCCTCATACTGTTCAATATAATGAATTCAAAGGAGGTGATAAAAATGGTTAGC |
| L23-2R | AGTATGAGGAAATCAAGAAGTAAAAAGAATAATGAAAGCATCCTAGTTACCCTCGAGATCTGG |
| L23-3F | ACTTGCATAAAAGTAGTGGTATTCATATTATTCAGTAAGTAAAAGGAGGTGATAAAAATGGTTAGC |
| L23-3R | TACCACTACTTTTATGCAAGTGACAATTCAAATTGAATTGCTAGTTACCCTCGAGATCTGG |
| L23-4F | ACTTGATTTATGCCAAAGAATGATGTATTATAATTACAAAAGGAGGTGATAAAAATGGTTAGC |
| L23-4R | TTCTTTGGCATAAATCAAGTGAATATTTGCGCTTGTATGCCACTAGTTACCCTCGAGATCTGG |
| L23-5F | TTTGGTTGAAATTGTCAATATCGCTTTGTAAAATCTGCGCGAAAGGAGGTGATAAAAATGGTTAGC |
| L23-5R | ATATTGACAATTTCAACCAAAAATTCAGGCAGTGGTCGTCCTAGTTACCCTCGAGATCTGG |
| L23-6F | TTTTAATTTGATAATTCATTGTGGTATTGCCTATATTTAAAGGAGGTGATAAAAATGGTTAGC |
| L23-6R | CAATGAATTATCAAATTAAAATTGCTAAAATAGTAAAAATAGTCTAGTTACCCTCGAGATCTGG |

| | |
|---|---|
| L23-7F | GATGTTGATATAAAAATCAACATCGTTATATTTTTGCGAAAGGAGGTGATAAAAATGGTTAGC |
| L23-7R | TGTTGATTTTTATATCAACATCGCAACATCACAAAAATGATGCACTAGTTACCCTCGAGATCTGG |
| L23-8F | TCCCTTTGAAGATATACCGATATGGACTATATTTGCAGCCGTAAAGGAGGTGATAAAAATGGTTAGC |
| L23-8R | ATCGGTATATCTTCAAAGGGAGAAATATGATTATATTACCTAGTTACCCTCGAGATCTGG |
| L23-9F | AATTGACCGGTTTTATCACTCTTATTATAAATCAGTCCAAAGGAGGTGATAAAAATGGTTAGC |
| L23-9R | TAAGAGTGATAAAACCGGTCAATTATATCACTCTTTATTATTTTCACTAGTTACCCTCGAGATCTGG |
| L23-10F | AGCGTAATTGATAAATACAATCATTTATTTATAATCTACGTTAAAAGGAGGTGATAAAAATGGTTAGC |
| L23-10R | ATTGTATTTATCAATTACGCTGTGAATAAATAAACTTTCTAGTTACCCTCGAGATCTGG |
| L23-11F | ACTTGATCAATTTACTCTGCACAATATATAATTACGCAAAGAAAAGGAGGTGATAAAAATGGTTAGC |
| L23-11R | GCAGAGTAAATTGATCAAGTATTGTTATTTAAATTGGTCTAGTTACCCTCGAGATCTGG |
| L23-12F | TGATAAGTGTAAGATAATTCACTATTATTAATACTGTCAAAGGAGGTGATAAAAATGGTTAGC |
| L23-12R | GTGAATTATCTTACACTTATCAATAACCTACAATAATATTAATACTAGTTACCCTCGAGATCTGG |
| L23-13F | TTTGAGTTATACTAATTCTACTTTAAAATGATTTAAAAGGAGGTGATAAAAATGGTTAGC |
| L23-13R | GTAGAATTAGTATAACTCAAAATAAAAGCCTGTGTTTTAAAACAGGCTAGTTACCCTCGAGATCTGG |
| L23-14F | TTGAAATTTAATCACGCCGGTCATATTATGTATTTATTAAAGGAGGTGATAAAAATGGTTAGC |
| L23-14R | TGACCGGCGTGATTAAATTTCAATGCGTCATGAAAAAGGATTCATCTAGTTACCCTCGAGATCTGG |
| L23-15F | TTACCGCCTTGATGTGTAAAATTAATTACGCTACTATGCGGTTAAAGGAGGTGATAAAAATGGTTAGC |
| L23-15R | AATTTTACACATCAAGGCGGTAATACTTTATAAATTGTGACTAGTTACCCTCGAGATCTGG |
| L23-16F | ACATTTTAGCAACCGCTATATATCATACTGCTATAATACTGATAAAGGAGGTGATAAAAATGGTTAGC |
| L23-16R | TATATAGCGGTTGCTAAAATGTTGTAAAGATAGAAATTTCTAGTTACCCTCGAGATCTGG |
| L23-17F | ATTGATTGGCAATATATTAAATTATATCATACCCCTTTAAAGGAGGTGATAAAAATGGTTAGC |
| L23-17R | TAATTTAATATATTGCCAATCAATCACTTATGTAAAGGATAAAAATCTAGTTACCCTCGAGATCTGG |
| L23-18F | TTGACTGGTAATTTGTGAATTATTGTGTTATAGTCCCGCCAAAAGGAGGTGATAAAAATGGTTAGC |
| L23-18R | AATTCACAAATTACCAGTCAATCCGGAAGACCGGCATTGTCTAGTTACCCTCGAGATCTGG |
| L23-19F | AGTTTTTGACAAATCGGCTCTGCTGGGCCGTATAATTGCCCCTAAAGGAGGTGATAAAAATGGTTAGC |
| L23-19R | GAGCCGATTTGTCAAAAACTCCATAATTCATCCGGCTCTAGTTACCCTCGAGATCTGG |

| | |
|---|---|
| L23-20F | AAATGCCTGTTGAAAATTTATATTAACATATAATATACTTGAAAGGAGGTGATAAAAATGGTTAGC |
| L23-20R | TATAAATTTTCAACAGGCATTTAATCCTTCATAGCGATTTACTAGTTACCCTCGAGATCTGG |
| EBI-19F | GACAAATCGGCTCTGCTGGGCCGTATAATTGCCCCTAAAGGAGGTGATAAAAATGACGGTCTGCGCAAA |
| EBI-19R | CAGCAGAGCCGATTTGTCAAAAACTCCATAATTCATCCGGCTCTCGAGGTGAAGACGAAAGG |
| EBI-10F | TTGATAAATACAATCATTTATTTATAATCTACGTTAAAAGGAGGTGATAAAAATGACGGTCTGCGCAAA |
| EBI-10R | TAAATGATTGTATTTATCAATTACGCTGTGAATAAATAAACTTTCTCGAGGTGAAGACGAAAGG |
| EBI-13F | TTTTGAGTTATACTAATTCTACTTTAAAATGATTTAAAAGGAGGTGATAAAAATGACGGTCTGCGCAAA |
| EBI-13R | GAATTAGTATAACTCAAAATAAAAGCCTGTGTTTTAAAACAGGCTCGAGGTGAAGACGAAAGG |
| EBI-11F | TCAATTTACTCTGCACAATATATAATTACGCAAAGAAAAGGAGGTGATAAAAATGACGGTCTGCGCAAA |
| EBI-11R | ATTGTGCAGAGTAAATTGATCAAGTATTGTTATTTAAATTGGTCTCGAGGTGAAGACGAAAGG |
| EBI-18F | TGGTAATTTGTGAATTATTGTGTTATAGTCCCGCCAAAAGGAGGTGATAAAAATGACGGTCTGCGCAAA |
| EBI-18R | CAATAATTCACAAATTACCAGTCAATCCGGAAGACCGGCATTGTCTCGAGGTGAAGACGAAAGG |
| EBI-5F | TTGAAATTGTCAATATCGCTTTGTAAAATCTGCGCGAAAGGAGGTGATAAAAATGACGGTCTGCGCAAA |
| EBI-5R | GCGATATTGACAATTTCAACCAAAAATTCAGGCAGTGGTCGTCCTCGAGGTGAAGACGAAAGG |
| EBI-4F | CTTGATTTATGCCAAAGAATGATGTATTATAATTACAAAAGGAGGTGATAAAAATGACGGTCTGCGCAAA |
| EBI-4R | TTCTTTGGCATAAATCAAGTGAATATTTGCGCTTGTATGCCACTCGAGGTGAAGACGAAAGG |

**Table S7. The strains used in this study.**

| Strains | Description | Source |
|---|---|---|
| *E. coli* JM109 | *recA1*, *endA1*, *thi*, *gyrA96*, *supE44*, *hsdR17*Δ (*lac-proAB*) /F'[*traD36,proAB*$^+$, *lacI*$^q$, *lacZ*Δ *M15*] | Laboratory stock |
| *B. subtilis* | Wild-type *Bacillus subtilis* 168 | Laboratory stock |
| BS-P43-sfGFP | Wild-type *B. subtilis* carrying the plasmid PXY-P43-sfGFP | Laboratory stock |
| BS-A1-sfGFP | Wild-type *B. subtilis* carrying the plasmid PXY-A1-sfGFP | This study |
| BS-A2-sfGFP | Wild-type *B. subtilis* carrying the plasmid PXY-A2-sfGFP | This study |
| BS-A3-sfGFP | Wild-type *B. subtilis* carrying the plasmid PXY-A3-sfGFP | This study |
| BS-A4-sfGFP | Wild-type *B. subtilis* carrying the plasmid PXY-A4-sfGFP | This study |
| BS-A5-sfGFP | Wild-type *B. subtilis* carrying the plasmid PXY-A5-sfGFP | This study |
| BS-A6-sfGFP | Wild-type *B. subtilis* carrying the plasmid PXY-A6-sfGFP | This study |
| BS-A7-sfGFP | Wild-type *B. subtilis* carrying the plasmid PXY-A7-sfGFP | This study |
| BS-A8-sfGFP | Wild-type *B. subtilis* carrying the plasmid PXY-A8-sfGFP | This study |
| BS-A9-sfGFP | Wild-type *B. subtilis* carrying the plasmid PXY-A9-sfGFP | This study |
| BS-A10-sfGFP | Wild-type *B. subtilis* carrying the plasmid PXY-A10-sfGFP | This study |
| BS-A11-sfGFP | Wild-type *B. subtilis* carrying the plasmid PXY-A11-sfGFP | This study |
| BS-A12-sfGFP | Wild-type *B. subtilis* carrying the plasmid PXY-A12-sfGFP | This study |
| BS-A13-sfGFP | Wild-type *B. subtilis* carrying the plasmid PXY-A13-sfGFP | This study |
| BS-A14-sfGFP | Wild-type *B. subtilis* carrying the plasmid PXY-A14-sfGFP | This study |
| BS-A15-sfGFP | Wild-type *B. subtilis* carrying the plasmid PXY-A15-sfGFP | This study |
| BS-A16-sfGFP | Wild-type *B. subtilis* carrying the plasmid PXY-A16-sfGFP | This study |
| BS-A17-sfGFP | Wild-type *B. subtilis* carrying the plasmid PXY-A17-sfGFP | This study |
| BS-A18-sfGFP | Wild-type *B. subtilis* carrying the plasmid PXY-A18-sfGFP | This study |
| BS-A19-sfGFP | Wild-type *B. subtilis* carrying the plasmid PXY-A19-sfGFP | This study |
| BS-A20-sfGFP | Wild-type *B. subtilis* carrying the plasmid PXY-A20-sfGFP | This study |
| BS-A21-sfGFP | Wild-type *B. subtilis* carrying the plasmid PXY-A21-sfGFP | This study |
| BS-A22-sfGFP | Wild-type *B. subtilis* carrying the plasmid PXY-A22-sfGFP | This study |
| BS-A23-sfGFP | Wild-type *B. subtilis* carrying the plasmid PXY-A1-sfGFP | This study |
| BS-A24-sfGFP | Wild-type *B. subtilis* carrying the plasmid PXY-A2-sfGFP | This study |
| BS-A25-sfGFP | Wild-type *B. subtilis* carrying the plasmid PXY-A3-sfGFP | This study |
| BS-A26-sfGFP | Wild-type *B. subtilis* carrying the plasmid PXY-A4-sfGFP | This study |
| BS-A27-sfGFP | Wild-type *B. subtilis* carrying the plasmid PXY-A5-sfGFP | This study |
| BS-A28-sfGFP | Wild-type *B. subtilis* carrying the plasmid PXY-A6-sfGFP | This study |
| BS-A29-sfGFP | Wild-type *B. subtilis* carrying the plasmid PXY-A7-sfGFP | This study |
| BS-A30-sfGFP | Wild-type *B. subtilis* carrying the plasmid PXY-A8-sfGFP | This study |
| BS-A31-sfGFP | Wild-type *B. subtilis* carrying the plasmid PXY-A9-sfGFP | This study |
| BS-A32-sfGFP | Wild-type *B. subtilis* carrying the plasmid PXY-A10-sfGFP | This study |
| BS-A33-sfGFP | Wild-type *B. subtilis* carrying the plasmid PXY-A11-sfGFP | This study |
| BS-A34-sfGFP | Wild-type *B. subtilis* carrying the plasmid PXY-A12-sfGFP | This study |
| BS-A35-sfGFP | Wild-type *B. subtilis* carrying the plasmid PXY-A13-sfGFP | This study |
| BS-A36-sfGFP | Wild-type *B. subtilis* carrying the plasmid PXY-A14-sfGFP | This study |
| BS-A37-sfGFP | Wild-type *B. subtilis* carrying the plasmid PXY-A15-sfGFP | This study |
| BS-A38-sfGFP | Wild-type *B. subtilis* carrying the plasmid PXY-A16-sfGFP | This study |
| BS-A39-sfGFP | Wild-type *B. subtilis* carrying the plasmid PXY-A17-sfGFP | This study |
| BS-A40-sfGFP | Wild-type *B. subtilis* carrying the plasmid PXY-A18-sfGFP | This study |
| BS-A41-sfGFP | Wild-type *B. subtilis* carrying the plasmid PXY-A19-sfGFP | This study |

| | | |
|---|---|---|
| BS-A42-sfGFP | Wild-type *B. subtilis* carrying the plasmid PXY-A20-sfGFP | This study |
| BS-A43-sfGFP | Wild-type *B. subtilis* carrying the plasmid PXY-A21-sfGFP | This study |
| BS-A44-sfGFP | Wild-type *B. subtilis* carrying the plasmid PXY-A12-sfGFP | This study |
| BS-A45-sfGFP | Wild-type *B. subtilis* carrying the plasmid PXY-A13-sfGFP | This study |
| BS-A46-sfGFP | Wild-type *B. subtilis* carrying the plasmid PXY-A14-sfGFP | This study |
| BS-A47-sfGFP | Wild-type *B. subtilis* carrying the plasmid PXY-A15-sfGFP | This study |
| BS-A48-sfGFP | Wild-type *B. subtilis* carrying the plasmid PXY-A16-sfGFP | This study |
| BS-A49-sfGFP | Wild-type *B. subtilis* carrying the plasmid PXY-A17-sfGFP | This study |
| BS-A50-sfGFP | Wild-type *B. subtilis* carrying the plasmid PXY-A18-sfGFP | This study |
| BS-A34-sfGFP | Wild-type *B. subtilis* carrying the plasmid PXY-A12-sfGFP | This study |
| JM109-J23119-sfGFP | Wild-type JM109 carrying the plasmid PXY- Ran3-sfGFP | This study |
| L23R7 | Wild-type L23 delete seven genes including *pta*, *adhE*, *plfB*, *frdBC*, *ldhA*, *fnr*, and *endA*. | Laboratory stock |
| L23R7-19 | The L23R7 overexpressed the *crtEBI* genes by the L23-19 promoter | This study |
| L23R7-10 | The L23R7 overexpressed the *crtEBI* genes by the L23-10 promoter | This study |
| L23R7-13 | The L23R7 overexpressed the *crtEBI* genes by the L23-13 promoter | This study |
| L23R7-11 | The L23R7 overexpressed the *crtEBI* genes by the L23-11 promoter | This study |
| L23R7-18 | The L23R7 overexpressed the *crtEBI* genes by the L23-18 promoter | This study |
| L23R7-5 | The L23R7 overexpressed the *crtEBI* genes by the L23-5 promoter | This study |
| L23R7-4 | The L23R7 overexpressed the *crtEBI* genes by the L23-4 promoter | This study |
